# Phenotypic and transcriptomic analyses reveal major differences between apple and pear scab nonhost resistance

**DOI:** 10.1101/2021.06.01.446506

**Authors:** Emilie Vergne, Elisabeth Chevreau, Elisa Ravon, Sylvain Gaillard, Sandra Pelletier, Muriel Bahut, Laure Perchepied

**Author notes:** Emilie Vergne and Elisabeth Chevreau made equal contributions to this work.

## Abstract

Nonhost resistance is the outcome of most plant/pathogen interactions, but it has rarely been described in Rosaceous fruit species. Apple (*Malus x domestica* Borkh.) have a nonhost resistance to *Venturia pyrina*, the scab species attacking European pear (*Pyrus communis* L.). Reciprocally, *P. communis* have a nonhost resistance to *Venturia inaequalis*, the scab species attacking apple. The major objective of our study was to compare the scab nonhost resistance in apple and in European pear, at the phenotypic and transcriptomic levels. Macro- and microscopic observations after reciprocal scab inoculations indicated that, after a similar germination step, nonhost apple/*V. pyrina* interaction remained nearly symptomless, whereas more hypersensitive reactions were observed during nonhost pear/*V. inaequalis* interaction. Comparative transcriptomic analyses of apple and pear nonhost interactions with *V. pyrina* and *V. inaequalis*, respectively, revealed differences. Very few differentially expressed genes were detected during apple/*V. pyrina* interaction, preventing the inferring of underlying molecular mechanisms. On the contrary, numerous genes were differentially expressed during pear/*V. inaequalis* interaction, allowing a deep deciphering. Pre-invasive defense, such as stomatal closure, could be inferred, as well as several post-invasive defense mechanisms (apoplastic reactive oxygen species accumulation, phytoalexin production and alterations of the epidermis composition). In addition, a comparative analysis between pear scab host and nonhost interactions indicated that, although specificities were observed, two major defense lines seems to be shared in these resistances: cell wall and cuticle potential modifications and phenylpropanoid pathway induction. This first deciphering of the molecular mechanisms underlying a nonhost scab resistance in pear offers new possibilities for the genetic engineering of sustainable scab resistance in this species. Concerning nonhost scab resistance in apple, further analyses must be considered with the aid of tools adapted to this resistance with very few cells engaged.

## Introduction

Apple (*Malus x domestica* Borkh.) and European pear (*Pyrus communis* L.) are two closely related species belonging to the Rosaceae family. Reclassification of the Rosaceae placed both *Pyrus* and *Malus* genera in the subfamily Spiraeoideae, tribe Pyreae and subtribe Pyrinae, this subtribe corresponding to the long-recognized subfamily Maloideae (Potter *et al*., 2007). Efforts to resolve relationships within this subtribe have frequently failed, and Campbell *et al*. (Campbell *et al*., 2007) concluded that the genera of this subtribe Pyreae have not diverged greatly genetically. The recent sequencing of the pear genome (Wu *et al*., 2013) allowed a precise comparison with the apple genome (Velasco *et al*., 2010) and led to the estimation of a divergence time between the two genera of ≈ 5.4 – 21.5 million years ago. Furthermore, apple and pear genomes share similar chromosome number (n=17), structure and organization.

Scab disease, caused by *Venturia* spp., affects several rosaceous fruit tree species. These hemibiotrophic pathogens can infect only a limited host-range during their parasitic stage, but they can overwinter as saprophytes in the leaf litter of a larger range of plant species (Stehmann *et al*., 2020). Scab disease is caused by *V. inaequalis* on apple, by *V. pyrina* (formerly named *V. pirina* (Rossman *et al*., 2018) on European pear, and by *V. nashicola* on Japanese (*P. pyrifolia* Nakai) and Chinese (*P. ussuriensis* Maxim) pears. Cross inoculations of *Venturia* spp. on different rosaceous fruit trees indicates that these pathogens are highly host specific, probably indicating a close co-evolution of these pathogens with their hosts (Gonzalez-Dominguez *et al*., 2017).

A plant species unable to be successfully infected by all isolates of a pathogen species is considered as a nonhost for this pathogen. Nonhost interactions of *Venturia* spp. on apple and pear have rarely been described. Microscopic observations have been made on *P. communis* / *V. nashicola* (Jiang *et al*., 2014) as well as *M. domestica* / *V. pyrina* and *P. communis* / *V. inaequalis* (Stehmann *et al*., 2020, Chevalier *et al*., 2004). In all cases, conidia germinated and produced appressoria and runner hypheae, but failed to establish a network of stroma. No macroscopic symptoms were visible.

Because of its durability, nonhost resistance has attracted numerous studies over the last decade, which have uncovered its multiple and complex defense components. The underlying mechanisms of nonhost resistance comprise pre-invasion resistance with preformed or induced cell-wall defenses, metabolic defense with phytoanticipin or phytoalexin accumulation, pattern-triggered immunity (PTI) as well as elicitor-triggered immunity (ETI) and various signaling pathways (Lee *et al*., 2017a).

To our knowledge, the molecular bases of scab nonhost resistance of apple and pear have never been investigated. We were not able to find reports on large-scale fungi nonhost resistance analyses in pear and the few available in apple are about Penicillium digitatum and are conducted on fruit (Buron-Moles *et al*., 2015, Vilanova *et al*., 2017).

However, it is possible to find genome-wide molecular analyses of scab host resistance in apple and pear. Thus, Perchepied *et al*. (Perchepied *et al*., 2021) performed a detailed transcriptomic analysis of the host resistance of pear against *V. pyrina* strain VP102, deployed in a transgenic pear bearing the well-known apple Rvi6 resistance gene against *V. inaequalis*. They reported the modulation of expression of 4170 genes and revealed that downstream of the pathogen recognition, the signal transduction was triggered with calcium, G-proteins and hormonal signaling (jasmonic acid (JA) and brassinosteroids), without involvement of salicylic acid (SA), and that this led to the induction of defense responses such as a remodeling of primary and secondary cell wall, cutin and cuticular waxes biosynthesis, systemic acquired resistance (SAR) signal perception in distal tissues, and the biosynthesis of phenylpropanoids (flavonoids and lignin). Only four other transcriptomic studies involving pear/pathogen host interactions have been published so far but are not concerning scab. Yan *et al*. (Yan *et al*., 2018) reported the modulation of expression of 144 pear genes after fruit treatment by Meyerozyma guilliermondii, an antagonistic yeast used for biocontrol of natural pear fruit decay. Zhang *et al*. (Zhang *et al*., 2020) similarly reported the modulation of expression of 1076 pear genes after treatment with Wickerhamomyces anomalus, another biocontrol agent. Using RNA-seq, Wang *et al*. (Wang *et al*., 2017a) reported a major role of ethylene (ET) signalization during the compatible interaction between P. pyrifolia and Alternaria alternata, a necrotrophic pathogen. Finally, Xu *et al*. (Xu *et al*., 2020) applied RNA-seq to characterize the genes of Penicillium expansum activated after infection of pear fruits.

Concerning host resistance of apple against *V. inaequalis*, subtractive hybridization (Paris *et al*., 2009, Cova *et al*., 2017) and cDNA-AFLP (Paris *et al*., 2012) led to the identification of a limited set of differentially expressed genes in Rvi6 natural resistant ‘Florina’ variety (scab inoculated ‘Florina’ versus mock, (Cova *et al*., 2017)), or in Rvi6 resistant transgenic ‘Gala’ lines (Rvi6 transgenic ‘Gala’ versus non-transformed ‘Gala’, after scab inoculation, (Paris *et al*., 2009); Rvi6 transgenic ‘Gala’ before versus post scab inoculation, (Paris *et al*., 2012)). Recently, Perchepied *et al*. (Perchepied *et al*., 2021) also performed a transcriptomic analysis of the Rvi6 resistance in a transgenic ‘Gala’ line (transgenic versus non-transformed, before and after scab inoculation). They reported the modulation of expression of 2977 genes and revealed that downstream of the pathogen recognition, signal transduction was triggered with calcium and interconnected hormonal signaling (auxins and brassinosteroids), without involvement of SA, and that this led to the induction of defense responses such as a remodeling of primary and secondary cell wall, galactolipids biosynthesis, SAR signal generation and the biosynthesis of flavonoids. Genome-wide molecular analyses of apple scab host resistance have also been achieved in other context than the Rvi6 resistance. A RNA-seq analyze identified five candidate genes putatively involved in the ontogenic scab resistance of apple (Gusberti *et al*., 2013). In addition, nuclear proteome analysis identified 13 proteins with differential expression patterns among varying scab resistance ‘Antonovka’ accessions (Sikorsskaite-Gudziuniene *et al*., 2017). Recently, Masoodi *et al*. (Masoodi *et al*., 2022) performed a RNA-seq analyze comparing three scab-resistant (‘Florina’, ‘Prima’, and ‘White Dotted Red’) and three susceptible (‘Ambri’, ‘Vista Bella’, and ‘Red Delicious’) apple genotypes out to mine new scab resistance genes. They reported the modulation of expression of 822 genes related to various pathways, i.e., metabolic, protein processing, biosynthesis of secondary metabolites, plant hormone signal transduction, autophagy, ubiquitin-mediated proteolysis, plant-pathogen interaction, lipid metabolism, and protein modification pathways.

Thus, if large-scale analyses of pear and apple scab host resistance can be found, in-depth knowledge of transcriptional patterns and gene functions involved in apple and pear scab nonhost resistance is still needed. The objectives of our study were 1) to precisely describe nonhost resistance symptoms in *M. domestica* / *V*.*pyrina* and *P. communis* / *V. inaequalis* interactions 2) to analyze the underlying molecular mechanisms of both nonhost interactions through a transcriptomic study 3) to compare the mechanism of host (Perchepied *et al*., 2021, Masoodi *et al*., 2022) and nonhost scab resistance in apple and European pear.

## Methods

### Biological material

Apple plants from the cultivar ‘Gala’ and pear plants from the cultivar ‘Conference’ were chosen because of their susceptibility to *V. inaequalis* and *V. pyrina*, respectively. The apple and pear genotypes were multiplied in vitro, rooted and acclimatized in greenhouse as described previously (Faize *et al*., 2003, Leblay *et al*., 1991).

For apple scab inoculation, the *V. inaequalis* monoconidial isolate used was EU-B05 from the European collection of *V. inaequalis* of the European project Durable Apple Resistance in Europe (Lespinasse *et al*., 2000). For pear scab inoculation, the monoconidial strain VP102 of *V. pyrina* was chosen for its aggressiveness on ‘Conference’ (Chevalier *et al*., 2008a).

### Scab inoculation procedure

Greenhouse growth conditions and mode of inoculum preparation were as described in Parisi and Lespinasse (Parisi *et al*., 1996) for apple and Chevalier *et al*. (Chevalier *et al*., 2008b) for pear. Briefly, the youngest leaf of actively growing shoots was tagged and the plants inoculated with a conidial suspension (2 × 105 conidia ml−1) of *Venturia* pyrina strain VP102 for apple and *Venturia* inaequalis strain EUB04 for pear. Symptoms were recorded at 14, 21, 28, 35 and 42 days after inoculation on 20 actively growing shoots (one shoot by plant) for each interaction. The type of symptoms was scored using the 6 class-scale of Chevalier *et al*. (Chevalier *et al*., 1991).

### Microscopic observations

Histological studies were made on samples stained with the fluorophore solophenylflavine (Hoch *et al*., 2005). In brief, leaf discs were rinsed in ethanol 50° before staining in a water solution of solophenylflavine 7GFE 500 (SIGMA-Aldrich, St Louis USA) 0.1% (v/v) for 10 min. The samples were first rinsed in deionized water, then in 25% glycerol for 10 min. Finally, the leaf samples were mounted on glass-slides in a few drops of 50% glycerol. They were examined with a wide-field epifluorescence microscope BH2-RFC Olympus (Hamburg, D) equipped with the following filter combination: excitation filter 395 nm and emission filter 504 nm.

### Transcriptomics experiment

Leaf samples were immediately frozen in liquid nitrogen and kept at -80°C until analysis. Sampling concerned the youngest expanded leaf of each plant labeled the day of the inoculation. Each biological repeat is a pool of three leaves from three different plants and two biological repeats (n=2) have been made by condition (genotype x treatment x time). Leaf samples taken just before inoculation (T0) and at 24 and 72 hours post inoculation (hpi), were then used to perform transcriptomics analyses.

For RNA extraction, frozen leaves were ground to a fine powder in a ball mill (MM301, Retsch, Hann, Germany). RNA was extracted with the kit NucleoSpin RNA Plant (Macherey Nagel, Düren, Germany) according to the manufacturer’s instructions but with a modification: 4% of PVP40 (4 g for 100 ml) were mixed with the initial lysis buffer RAP before use. Purity and concentration of the samples were assayed with a Nanodrop spectrophotometer ND-1000 (ThermoFisher Scientific, Waltham, MA, USA) and by visualization on agarose gel (1% (weight/volume) agarose, TAE 0.5x, 3% (volume/volume) Midori green). Intron-spanning primers (forward primer: CTCTTGGTGTCAGGCAAATG, reverse primer: TCAAGGTTGGTGGACCTCTC) designed on the EF-1α gene (accession AJ223969 for apple and PCP017051 for pear, available at https://www.rosaceae.org/, with the datasets on “Pyrus communis v1.0 draft genome”) were used to check the absence of genomic DNA contamination by PCR. The PCR reaction conditions were as follows: 95°C for 5 min, followed by 35 cycles at 95°C for 30 s, 60°C for 45 s, 72°C for 1 min, with a final extension at 72°C for 5 min. The PCR products were separated on a 2% agarose gel.

Amplifications (aRNAs) were produced with MessageAmpII aRNA Kit (Ambion Invitrogen, Waltham, MA, USA), from 300 ng total RNA. Then 5 µg of each aRNA were retrotranscribed and labelled using a SuperScript II reverse transcriptase (Transcriptase inverse SuperScript™ II kit, Invitrogen, Carlsbad, CA, USA) and fluorescent dyes: either cyanine-3 (Cy3) or cyanine-5 (Cy5) (Interchim, Montluçon, France). Labeled samples (30 pmol each, one with Cy3, the other with Cy5) were combined two by two, depending on the experimental design (i. e. for example, the CF/EUB05/24 hpi sample and the CF/non inoculated sample are labeled with Cy3 and Cy5 respectively and pooled to co-hybridize the microarray). For each comparison two biological replicates were analyzed in dye-switch as described in Depuydt *et al*. (Depuydt *et al*., 2009). Paired labeled samples were then co-hybridized to Agilent microarray AryANE v2.0 (Agilent-070158_IRHS_AryANE-Venise, GPL26767 at GEO: https://www.ncbi.nlm.nih.gov/geo/) for apple, or Pyrus v1.0 (Agilent-078635_IRHS_Pyrus, GPL26768 at GEO) for pear, containing respectively 133584 (66792 sense and 66792 anti-sense probes) and 87812 (43906 sense and 43906 anti-sense probes) 60-mer oligonucleotide probes. The hybridizations were performed as described in Celton, Gaillard *et al*. (Celton *et al*., 2014) using a MS 200 microarray scanner (NimbleGen Roche, Madison, WI, USA).

For microarray analysis we designed two new chips. For apple we used a deduplicated probeset from the AryANE v1.0 ((Celton *et al*., 2014); 118740 probes with 59370 in sense and 59370 in anti-sense) augmented by 14844 probes (7422 in sense and 7422 in anti-sense) designed on new gene annotations from *Malus x domestica* GDDH13 v1.1 (https://iris.angers.inra.fr/gddh13 or https://www.rosaceae.org/species/malus/malus_x_domestica/genome_GDDH13_v1.1). These probes target new coding genes with UTRs when available, manually curated micro-RNA precursors and transposable elements. For transposable elements we used one consensus sequence for each family and a randomly peaked number of elements proportional to their respective abundance in the genome. The microarrays used in this study also have probes for coding genes of *V. inaequalis* but they have not been taken into account.

For pear the design was done on the *Pyrus communis* Genome v1.0 Draft Assembly & Annotation available on GDR (https://www.rosaceae.org/species/pyrus/pyrus_communis/genome_v1.0) web site. We have downloaded the reference genome and gene predictions fasta files and structural annotation gff file the 21st of September 2015. Using home-made Biopython scripts we have extracted spliced CDS sequences with 60 nucleotides before start and after stop codons to get UTR-like sequences likely to be found on transcripts resulting in a fasta file containing 44491 sequences. These 60 nucleotides increase the probability of finding specific probes on genes with high similarity. This file was sent to the eArray Agilent probe design tool (https://earray.chem.agilent.com/earray/) to generate one probe per gene prediction. Options used were: Probe Length: 60, Probe per Target: 1, Probe Orientation: Sense, Design Options: Best Probe Methodology, Design with 3’ Bias. The probeset was then reverse-complemented to generate anti-sense probes and filtered to remove duplicated probes. The final probeset contains 87812 unique probes targeting 1 (73612 probes) or more (14200 probes) potential transcript both in sense and anti-sense.

Normalization and statistical analyses performed to get normalized intensity values have been done as in Celton, Gaillard *et al*. (Celton *et al*., 2014). Briefly, data were normalized with the lowess method, and differential expression analyses were performed using the lmFit function and the Bayes moderated t-test with the LIMMA package in R software (Smyth, 2005, Ritchie *et al*., 2015). The pipeline AnaDiff used for these differential analyses is available at https://doi.org/10.5281/zenodo.6477917 (Pelletier, 2022). For each comparison and each probe, we retrieved a ratio of the logarithms of the fluorescence intensities (one per compared sample: T0 versus 24 hpi or T0 versus 72 hpi in our case) and an associated p-value. The applied p-value threshold to determine DEGs (differentially expressed genes) was 0.01. Through blast analysis, a TAIR accession number (The Arabidopsis Information Resource; https://www.arabidopsis.org/; (Berardini *et al*., 2015) has been linked to a majority of apple or pear “probe/corresponding gene”. Thanks to the Functional Classification SuperViewer tool (Provart *et al*., 2003), the TAIR accessions have then been used to class DEGs in functional categories according to MapMan software (https://mapman.gabipd.org/homemapman.gabipd.org; file Ath_AGI_LOCUS_TAIR10_Aug2012.txt; (Thimm *et al*., 2004)), and to highlight the enriched ones by calculating a normed frequency of the relative proportion of each category in DEGs set (input set) and reference set (Arabidopsis genes in MapMan database), and bootstrapping the dataset to provide a confidence estimate for the accuracy of the result. Normed Frequency is calculated as follows: (Number_in_Category_input_set_/Number_Classified_input_set_)/(Number_in_Category_reference_set_/Number_Classifi ed_reference_set_).

The detailed analysis of DEGs has been done through TAIR and KEGG (https://www.genome.jp/kegg/) databases, and bibliography. Metadata for the 193 (184 for pear, corresponding to 158 different functions and 9 for apple) DEGs discussed in this work are available in Table S1 and S2 (Online only).

### qPCR validation of transcriptomic data

In order to validate transcriptomic data, qPCR was performed on a selection of gene/sample associations. First-strand cDNA was synthesized using total RNA (2.0 μg) in a volume of 30 μl of 5× buffer, 0.5 μg of oligodT15 primer, 5 μl of dNTPs (2.5 mM each), and 150 units of MMLV RTase (Promega, Madison, WI, USA). The mixture was incubated at 42°C for 75 min. qPCR was then performed. Briefly, 2.5 µl of the appropriately diluted samples (1/16 dilution) were mixed with 5 µl of PerfeCTa SYBR Green SuperMix for iQ kit (Quantabio, Beverly, MA, USA) and 0.2 or 0.6 µl of each primer (10 µM) in a final volume of 10 µl. Primers were designed with Primer3Plus, their volumes were according to their optimal concentration (determined for reaction efficiency near to 100%; calculated as the slope of a standard dilution curve; (Pfaffl, 2001). Accessions, primer sequences and optimal concentrations are indicated in Table S3. The reaction was performed on a CFX Connect Real-Time System (BIO-RAD, Hercules, CA, USA) using the following program: 95°C, 5 min followed by 40 cycles comprising 95°C for 3 s, 60°C for 1 min. Melting curves were performed at the end of each run to check the absence of primer-dimers and nonspecific amplification products. Expression levels were calculated using the ΔΔCT method (Livak *et al*., 2001) and were corrected as recommended in Vandesompele *et al*. (Vandesompele *et al*., 2002), with three internal reference genes (GADPH, TUA and ACTIN 7 for apple, GADPH, TUA and EF1α for pear) used for the calculation of a normalization factor. For each couple DEG/sample (sample defining a plant, time, treatment and biological repeat combination), the ratio was obtained by dividing the mean value of CT calculated from 3 technical repeats by the normalization factor obtained for this sample.

## Results

### Macroscopic and microscopic symptom analysis

Nonhost interactions were observed in a test performed on leaves of ‘Gala’ apple and ‘Conference’ pear cultivars, inoculated by a *V. pyrina* strain (VP102) and a *V. inaequalis* strain (VI EUB05) respectively. At the macroscopic level, a total absence of sporulation was observed on all nonhost interactions (Table 1), on the contrary to host interactions (Fig. 1 A and B). Very few pear plants inoculated with *V. inaequalis* presented resistance symptoms such as pin points (Fig. 1C) and chlorotic lesions (Fig. 1D), whereas the apple ‘Gala’ remained completely symptomless after *V. pyrina* inoculations (Fig. 1E).

**Table 1:** Scab qualitative note of pear and apple lines inoculated with *V. pyrina* and *V. inaequalis*

| Percentage (number) of plants in the different classes of symptoms, 42 days after inoculation |  |  |  |  |
| --- | --- | --- | --- | --- |
| Class of symptoms | *V. pyrina* strain VP102 |  | *V. inaequalis* strain EUB05 |  |
|  | 'Conference' | 'Gala' | 'Conference' | 'Gala' |
| 0: absence of symptoms | 0 | 100 (28) | 90 (17) | 0 |
| 1: hypersensitivity (pin points) | 0 | 0 | 5 (1) | 0 |
| 2: resistance (chlorotic lesions, slight necrosis, crinkled aspect) | 0 | 0 | 5 (1) | 0 |
| 3a: weak resistance (necrotic or chlorotic lesions with occasional very light sporulation) | 0 | 0 | 0 | 0 |
| 3b: weak susceptibility (clearly sporulating chlorotic or necrotic lesions) | 0 | 0 | 0 | 0 |
| 4: susceptibility (sporulation only) | 100 (19) | 0 | 0 | 100 (28) |

**Figure 1:**
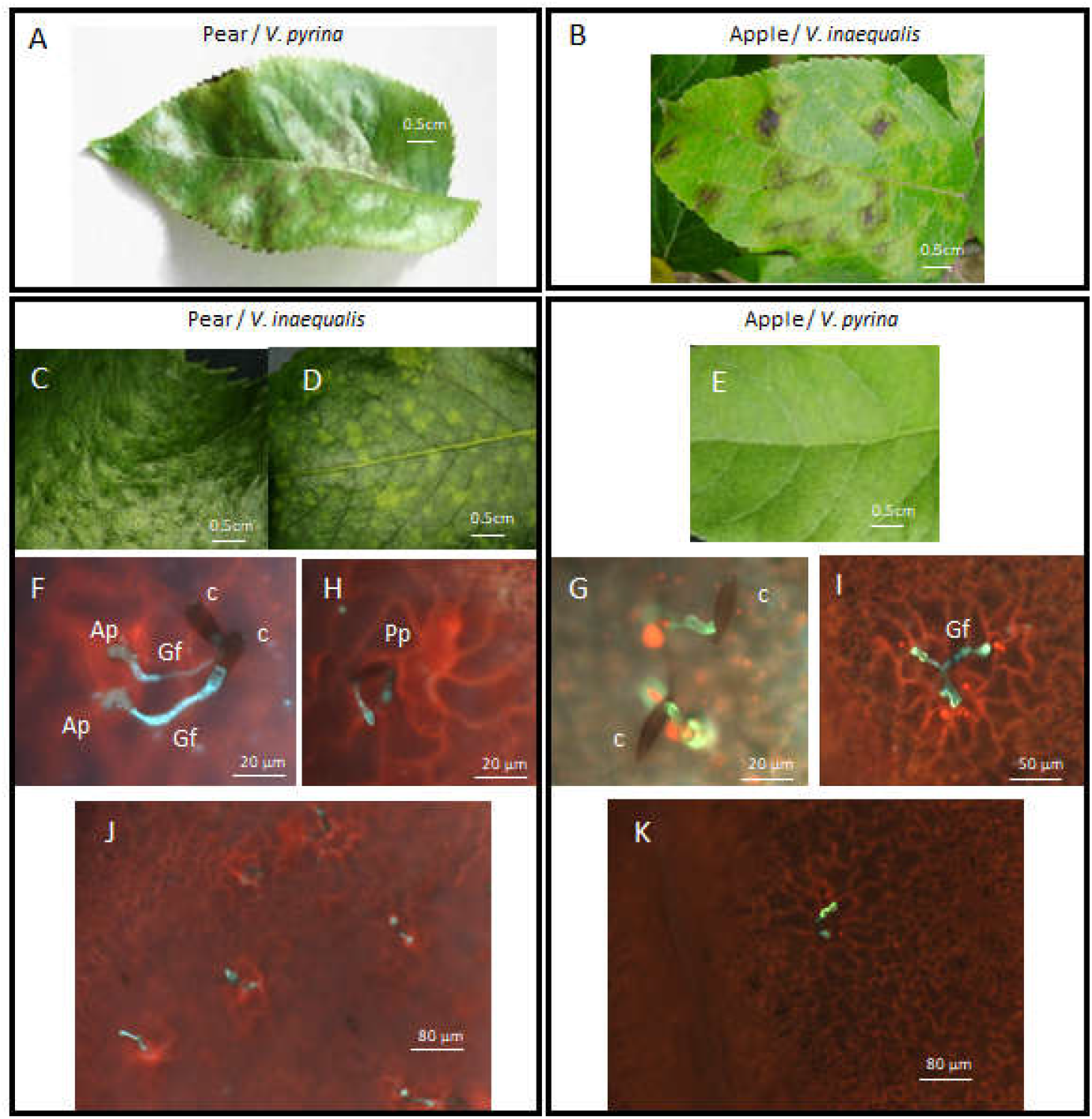
Macro- and microscopic observations of nonhost interactions. *V. pyrina* VP102 strain / pear ‘Conference’ (A) and *V. inaequalis* EUB05 strain / apple ‘Gala’ (B) 21 days symptoms are shown as classical ones of susceptible host interactions, to compare to nonhost ones. Binocular observation 21 days after *V. inaequalis* EUB05 strain inoculation on ‘Conference’ (C) and (D) and *V. pyrina* VP102 strain inoculation on ‘Gala’ (E). Wide field fluorescence observations of: ‘Conference’ 3 days (F) and 14 days (H and J) after *V. inaequalis* EUB05 strain inoculation, ‘Gala’ 3 days (G) and 14 days (I and K) after *V. pyrina* VP102 strain inoculation. Ap: appressorium, C: conidia, Gf: germination filament, Pp: pin point

At the microscopic level, three days after inoculation, there was no clear difference between host and nonhost interactions: the conidia of *V. inaequalis* and *V. pyrina* germinated equally on both hosts forming one or two appressoria (Fig. 1 F and G). However, 14 days after inoculation, there was a clear reaction of the plant cells in contact with the appressoria (accumulation of red autofluorescent compounds and enlargement of these cells), which could indicate very small scale hypersensitive reactions (HR) (Fig. 1 H and I) in both plant species, more frequently in pear than in apple (Fig1. J and K). No formation of subcuticular stroma and no conidiogenesis were observed in the nonhost interactions.

### Global gene expression analysis

DEGs were obtained by comparing transcript abundance in leaves between T0 and 24 hpi and between T0 and 72 hpi, in the nonhost interactions ‘Gala’ / *V. pyrina* VP102 and ‘Conference’ / *V. inaequalis* EUB05. These time points were chosen in order to cover the period of establishment of the first intimate contacts between fungal and plant cells: conidia germination and appressoria formation. For each comparison, the experimental design is a dye switch approach (Mary-Huard *et al*., 2008) between the two biological repeats made by condition (genotype x treatment x time). Each biological repeat is a pool of three leaves from three different plants.

In total, 60 DEGs in apple and 1857 DEGs in pear were identified, which amounts to 0.19 % of all apple genes on the apple AryANE v2.0 microarray, and 4.23 % of all pear genes on the Pyrus v1.0 microarray (Table 2). Among the 1857 pear DEGs, 80.2 % were only detected at 24 hpi and 15.4 % only at 72 hpi, whereas 4.2 % were up-regulated or down-regulated similarly at both time points of the experiment. Among all the pear DEGs observed at 24 and 72 hpi, the proportion of up-regulated DEGs was higher (68.8 %) than the proportion of down-regulated DEGs (31.2 %).

**Table 2:** Number of DEGs identified during apple and pear nonhost response to *V. pyrina* and *V. inaequalis*

|  | 'Gala' / VP102 |  | 'Conference' / EUB05 |  |
| --- | --- | --- | --- | --- |
|  | 24 hpi | 72 hpi | 24 hpi | 72 hpi |
| Total # of DEGs* | 49 | 11 | 1570 | 364 |
| DEGs in % of all genes on the microarray** | 0.16 | 0.03 | 3.58 | 0.83 |
| % of DEGs up-regulated | 67.3 | 36.4 | 74.5 | 25.5 |
| % of DEGs down-regulated | 32.7 | 63.6 | 25.5 | 74.5 |
| % of DEGs without *Arabidopsis* homolog | 27.1 | 30.4 | 0.70 | 1.09 |

Note: \*: DEGs numbers were calculated using the p-adj values $\leq 0.01$ as selection threshold, \*\*: 31311 genes on the apple Ariane V2 microarray, 43906 genes on the pear V1 microarray

To validate the transcriptomic data, 12 DEGs with varied ratios (between -1.9 and 2.9) were tested by quantitative RT-PCR (qPCR; Table S3), on the two biological repeats used for transcriptomic analyses. Considering the low number of DEGs found for apple in this study, we only tested two of them in qPCR. As seen in Table 2 for pear, at 24 hpi, a majority of DEGs are up-regulated and at 72 hpi, a majority of DEG are down-regulated. qPCR was then performed essentially on DEGs with positive ratios at 24 hpi and negative ratios at 72 hpi (Table S3). The qPCR results confirmed the induced or repressed status of all tested DEGs.

Among the 1857 pear DEGs, the 1845 DEGs with Arabidopsis homologs (TAIR identifier) have been classified according to MapMan functional categories (Fig. 2). In order to highlight the enriched classes, a normed frequency of the relative proportion of each category in DEGs set and reference set (Arabidopsis genes in MapMan database) has been calculated and bootstraps have been done to provide a confidence estimate for the accuracy of the output (Fig. 2). We then more particularly explored the DEGs present in the following enriched classes: Hormone metabolism, Stress, Lipid metabolism, Signaling, Secondary metabolism, Cell wall, and depending on the defense pathways and responses identified, search for others related DEGs in enriched wider classes: Protein, RNA and Miscellaneous. The Table 3 gives the defense pathways and responses identified and the functional categories in which DEGs related to these pathways/responses have been found. The analyze of the 183 DEGs found (corresponding to 157 different functions) is expanded in the Discussion section (Metadata of the 183 DEGs in Table S1).

**Table 3:**
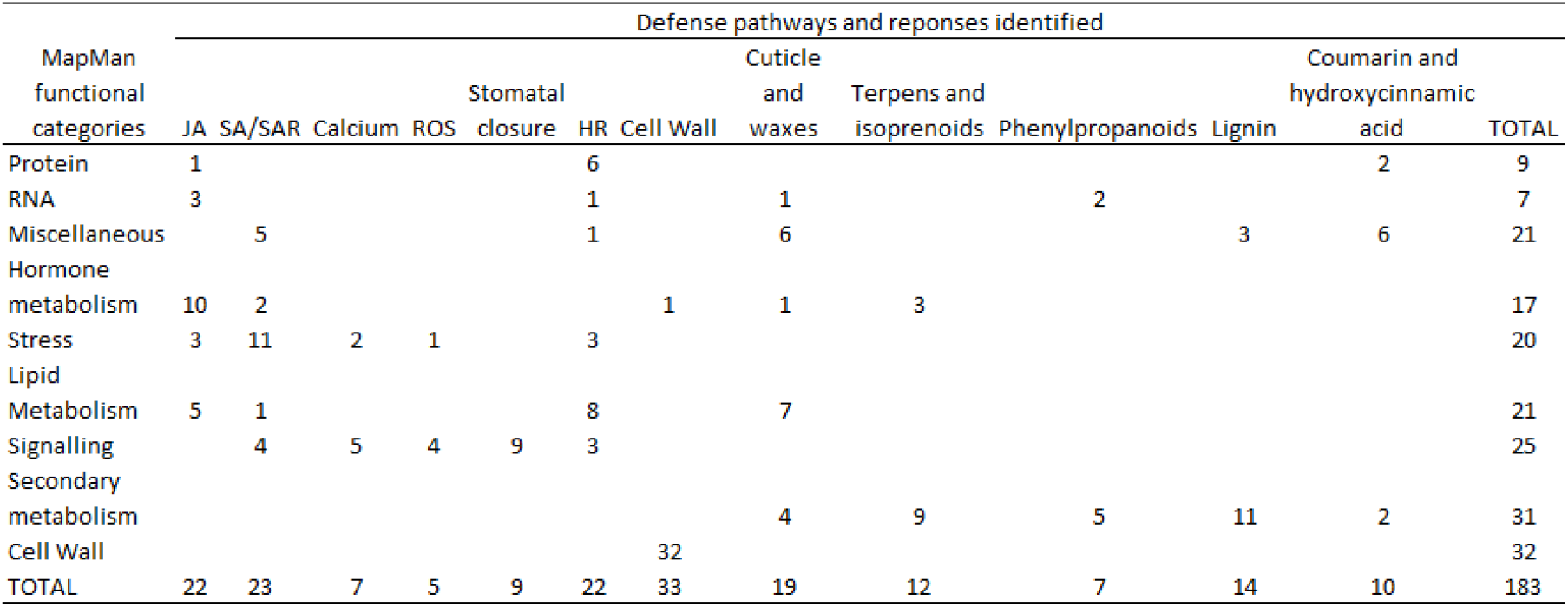
Numbers of DEGs in the defense pathways and responses identified among the enriched functional categories analyzed in pear nonhost response to *V. inaequalis*

| MapMan functional categories | Defense pathways and responses identified |  |  |  |  |  |  |  |  |  |  |  | TOTAL |
| --- | --- | --- | --- | --- | --- | --- | --- | --- | --- | --- | --- | --- | --- |
|  | JA | SA/SAR | Calcium | ROS | Stomatal closure | HR | Cell Wall | Cuticle and waxes | Terpens and isoprenoids | Phenylpropanoids | Lignin | Coumarin and hydroxycinnamic acid |  |
| Protein | 1 |  |  |  |  | 6 |  |  |  |  |  | 2 | 9 |
| RNA | 3 |  |  |  |  | 1 |  | 1 |  | 2 |  |  | 7 |
| Miscellaneous |  | 5 |  |  |  | 1 |  | 6 |  |  | 3 | 6 | 21 |
| Hormone metabolism | 10 | 2 |  |  |  |  | 1 | 1 | 3 |  |  |  | 17 |
| Stress | 3 | 11 | 2 | 1 |  | 3 |  |  |  |  |  |  | 20 |
| Lipid |  |  |  |  |  |  |  |  |  |  |  |  |  |
| Metabolism | 5 | 1 |  |  |  | 8 |  | 7 |  |  |  |  | 21 |
| Signalling |  | 4 | 5 | 4 | 9 | 3 |  |  |  |  |  |  | 25 |
| Secondary metabolism |  |  |  |  |  |  |  | 4 | 9 | 5 | 11 | 2 | 31 |
| Cell Wall |  |  |  |  |  |  | 32 |  |  |  |  |  | 32 |
| TOTAL | 22 | 23 | 7 | 5 | 9 | 22 | 33 | 19 | 12 | 7 | 14 | 10 | 183 |

**Figure 2:**
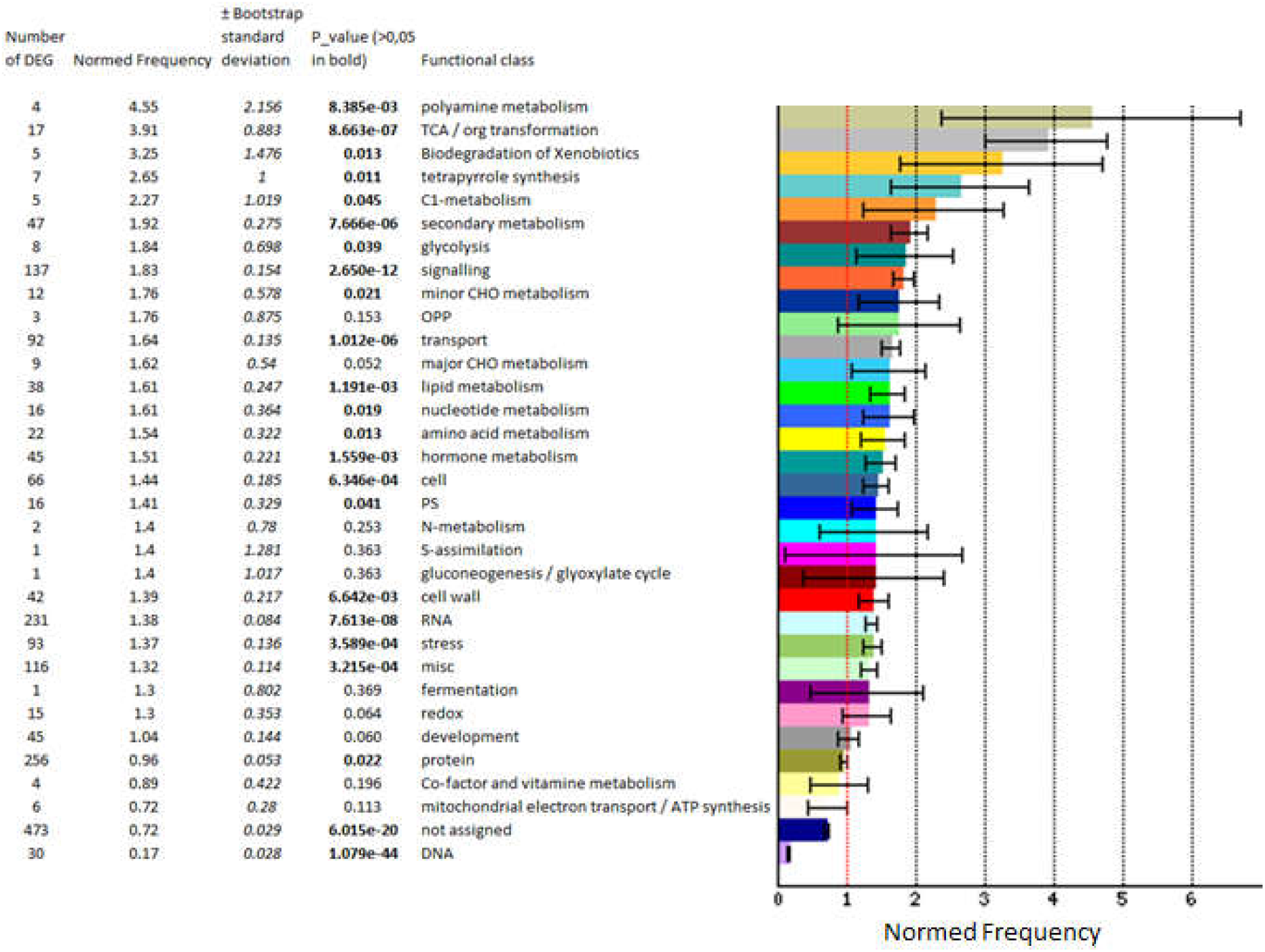
The 1845 DEGs with TAIR names have been classified according to MapMan functional categories. In order to highlight the enriched ones, a normed frequency of the relative proportion of each category in DEGs set (input set) and reference set (Arabidopsis genes in MapMan database) has been calculated as follows: Number_in_Category_input_set_/Number_Classified_input_set_)/(Number_in_Category_reference_set_/Number_Clas sified_reference_set_) and bootstraps have been done to provide a confidence estimate for the accuracy of the output

Among the 60 apple DEGs found, 17 have no Arabidopsis homolog, 12 more have unpredicted function and for 11 more we could not find information (Metadata of the 60 DEGs in Table S4). In view of our findings in pear / *V. inaequalis* nonhost interaction, 9 of the 20-remaining apple DEGS could be relevant in apple / *V. pyrina* nonhost interaction and are further analyzed in the Discussion section (Metadata of the 9 DEGs in Table S2).

## Discussion

### More frequent symptoms of nonhost resistance in pear versus apple at microscopic level

The apple ‘Gala’ remained completely symptomless after *V. pyrina* inoculations (Fig. 1E). This is similar to the observation of Chevalier *et al*. (Chevalier *et al*., 2004) after inoculation of ‘Gala’ with another *V. pyrina* strain. On the contrary, pear plants inoculated with *V. inaequalis* presented occasional pin points symptoms (Fig. 1D) and chlorotic lesions (Fig. 1E). Chlorotic lesions had already been observed by Chevalier *et al*. (Chevalier *et al*., 2004) after inoculation of the pear ‘Pierre Corneille’ with the *V. inaequalis* strain EUB04, but pin points had never been reported in this nonhost interaction.

At the microscopic level, 14 days after inoculation, there was a clear reaction of the plant cells in contact with the appressoria (accumulation of red autofluorescent compounds and enlargement of these cells), which could indicate very small scale hypersensitive reactions (HR) (Fig. 1 H and I, I and K) in both plant species. No formation of subcuticular stroma and no conidiogenesis were observed in the nonhost interactions, contrary to the host-resistance reactions (Perchepied *et al*., 2021). These observations are similar to the collapsed cells described by Chevalier *et al*. (Chevalier *et al*., 2004) in apple and pear nonhost reactions, and to the rare HR-like reactions observed by Stehmann *et al*. (Stehmann *et al*., 2001) on apple inoculated by *V. pyrina*.

Our results seem to indicate that the leaf surface morphology of apple and pear is equally compatible with *V. pyrina* and *V. inaequalis* conidia germination, without specific inhibition at this stage. Recognition probably occurs only at the appressorium site, leading to the cellular reactions observed. These reactions were limited to a few cells without visible symptoms in apple / *V. pyrina* interaction, but more extended in pear / *V. inaequalis* interaction and could occasionally produce macroscopic symptoms.

### Different patterns of global gene expression in nonhost resistance in pear versus apple

DEGs were analyzed by comparing transcript abundance in leaves between T0 and 24 hpi and between T0 and 72 hpi, in the nonhost interactions ‘Gala’ / *V. pyrina* VP102 and ‘Conference’ / *V. inaequalis* EUB05. This experimental design is open to criticism because it could include in results DEGs responding to the inoculation method (spray of water), and DEGs which expression varies due to leaves ageing. Water spray is effectively perceived as a mechanical stimulus by plants, the Arabidopsis response being largely regulated by the JA pathway induction under the control of MYC2/MYC3/MYC4 transcription factors (Van Moerkercke *et al*., 2019). But this response is transient and really fast as most of genes differentially regulated peak within 30 minutes regain untreated transcriptional levels within 3 hours (Van Moerkercke *et al*., 2019). It seems therefore very unlikely that our analysis at 24 and 72 hpi includes genes responding to water spray. Resistance due to leaves ageing i.e. ontogenic resistance has been investigated in *Malus*-*Venturia* pathosystem at 72 and 96 hpi, and 5 genes have been identified whose modulation could be linked to this resistance (Gusberti *et al*., 2013). None of these apple genes, or homologs in pear, were found differentially expressed at 72 hpi in our interactions (Table S5), which argue against the presence of ageing responding genes in our results. In total, 60 DEGs in apple and 1857 DEGs in pear were identified, which amounts to 0.19 % of all apple genes on the apple AryANE v2.0 microarray, and 4.23 % of all pear genes on the Pyrus v1.0 microarray (Table 2).

The very small number of DEGs (60) detected in the apple/ *V. pyrina* nonhost interaction at 24 or 72 hpi seems in agreement with the total absence of macroscopic symptoms observed during this interaction, and the few small HR-like reactions detected at the microscopic level 14 days post inoculation. Because these reactions involve only a few cells in the leaves, the changes in gene expression are probably below the threshold of DEG detection applied in this experiment. It is also possible that the mild response of apple to *V. pyrina* occurs later than 72 hpi, between 72 hpi and 6 days after inoculation. Indeed, Chevalier *et al*. (Chevalier *et al*., 2004) observed these rare HR from 6 days post inoculation, with no more evolution until 14 days after inoculation. A later time (6 days post inoculation) would have been necessary to conclude on this last hypothesis.

On the contrary to apple/ *V. pyrina* nonhost interaction, the number of DEGs (1857) detected during the pear / *V. inaequalis* interaction is in the same order of magnitude as the number of DEGs detected during pear host resistance to *V. pyrina* (see (Perchepied *et al*., 2021). This could be consistent with the more frequent observation in this interaction of microscopic HR-like reactions detected 14 days post inoculation, and of occasional macroscopic symptoms of resistance (chlorotic lesions or pin points).

### Hormone signaling pathways classically associated to resistance seems weakly involved in pear / *V. inaequalis* interaction

We found pear DEGs that seems to indicate that the JA pathway was repressed. The JA biosynthesis and metabolic conversions were reviewed by Wasternack *et al*. (Wasternack *et al*., 2018). In our data, at 24 hpi, the first step of JA biosynthesis, corresponding to the conversion of linoleic acid in 12-oxo-phytodienoic acid (OPDA), seems to be compromised: six out of seven lipoxygenases (LOX) (three LOX1, two LOX2 and two LOX5) are repressed, the last one being induced (Fig. 3). OPDA produced in the chloroplast is then transported to the peroxisome for subsequent conversion to JA via the action of OPR3 (12-oxo-phytodienoic acid reductase) and β-oxidation enzymes (reviewed in (Li *et al*., 2005) and in (Wasternack *et al*., 2018)). In pear, three β-oxidation enzymes were found activated more or less rapidly: ACX4 (24 hpi), MFP2 (72 hpi) and the thioesterase homolog to At2g29590 (72 hpi), which could suggest that constitutive OPDA stocks were turned into JA. But the early and long-lasting induction of JMT and ST2A genes could be in favor of a rapid conversion of JA in inactive compounds.

**Figure 3:**
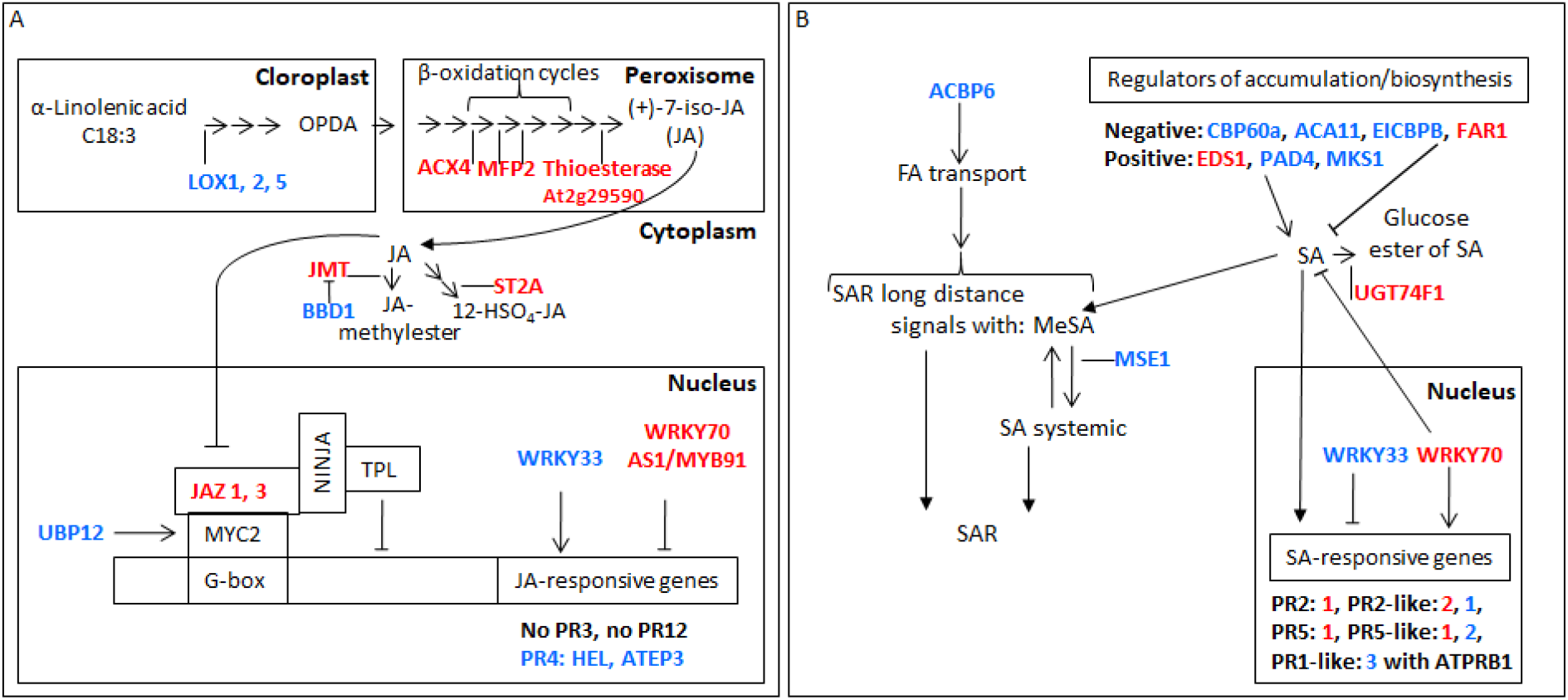
DEGs involved in hormonal pathways during pear/*V. inaequalis* nonhost interaction. A: DEGs involved in JA pathway; B: DEGs involved in SA pathway. Genes written in red are induced, genes written in blue are repressed. ACA11: autoinhibited Ca2+-ATPase, calmodulin-activated Ca2+ pumps at the plasma membrane, endoplasmic reticulum (ER), and vacuole. ACBP6: acyl-CoA-binding protein. ACX4: acyl-CoA-oxidase1. AS1/MYB91: Asymmetric leaves 1 transcription factor, CAMTA1: calmodulin-binding transcription activator, CBP60a: calmodulin-binding protein 60a, EDS1: enhanced disease suceptibility 1. FAR1: FAR-red impaired response 1. G-box: cis-element in the promoter. JAZ: jasmonate-zim domain protein, JMT: jasmonic acid carboxyl methyltransferase. LOX: lipoxygenase, MES1: methylesterase 1. MFP2: multifunctional protein 2. MKS1: MAP kinase substrate 1. MYC2: transcription factor. NINJA: novel interactor of JAZ. PAD4: phytoalexin deficient 4. UGT74F1: glucosyltransferase. PR1-like (with ATPRB1), PR2, PR3, PR4 (HEL and ATEP3), PR5, PR12: pathogenesis-related proteins. ST2A: sulfotransferase 2A. TPL: TOPLESS co-repressor. UBP12: ubiquitin-specific protease 12. WRKY: transcription factor

The behavior of some DEGs seemed indicate that the defense response depending on JA was also repressed in pear (Fig. 3). The transcription activator MYC2 of JA-induced genes is known to be repressed by its interaction with JAZ proteins (reviewed in (Wasternack *et al*., 2018)), and two JAZ1 and one JAZ3 coding genes were found activated at 24 hpi in pear. UBP12 is known as a stabilizer of MYC2 (Jeong *et al*., 2017). In our data, UBP12 was found repressed at 72 hpi, which could reinforce the inactivation of MYC2. WRKY33 is known as an activator of the JA defense pathway (Birkenbihl *et al*., 2012) and WRK70 (Kaurilind *et al*., 2015) or AS1 (or MYB91; (Nurmberg *et al*, 2007)) as inhibitors, and among JA-responsive proteins, the pathogenesis-related PR3, PR4 and PR12 act downstream MYC2 activation (Ali *et al*., 2018). In our data, accordingly with the repression of the activator WRKY33 and the activation of the inhibitors WRK70 and AS1, some JA-responsive genes were also found repressed, such as the chitinase coding genes PR4 (also called HEL) and ATEP3. Furthermore, no DEGs were found for PR3 and PR12 functions. To conclude, in the nonhost interaction between pear and *V. inaequalis*, some JA could be produced, but is rapidly converted into inactive compounds and the subsequent defense response seems to be repressed.

Pear DEGs were found that could indicate that the SA pathway was slightly engaged and rapidly repressed (Fig. 3). WRKY70 was induced at 72 hpi in our data. This transcription factor is known as a negative regulator of SA biosynthesis but a positive regulator of SA-mediated defense genes in Arabidopsis (Li *et al*., 2004, Li *et al*., 2006, Wang *et al*., 2006), among them PR2, PR5 but not PR1 (Li *et al*., 2017). WRKY33 which is known as a negative regulator of SA-responsive genes (Genot *et al*., 2017), was also repressed at 72 hpi in our data. PR2 and 5 are well-known anti-fungal proteins Hu *et al*., 1997, Mestre *et al*., 2017, Zhang *et al*., 2019). At 24 hpi a PR2, two PR2-like, a PR5 and a PR5-like coding genes were found induced in our work, another PR2-like and two others PR5-like being repressed. The differential expression was maintained at 72 hpi for only two of the previously activated ones. Furthermore no DEG was found for the PR1 function but three PR1-like genes were found repressed: ATPRB1 and genes homolog to At5g57625 and At4g33720. ATPRB1 was already reported as repressed by SA treatment (Santamaria *et al*., 2001). In our data, the WRKY70 transcription factor was induced later than the induced PR genes so we could imagine that induced PR genes were activated by another precocious regulation, such as an oxidative burst (see below), rather than by WRKY70. Furthermore, WRKY70 induction seems not sufficient to enable a long lasting induction of these defense genes.

Other pear DEGs seems to indicate that SA accumulation was transient. CBP60a (Truman *et al*., 2013), ACA11 (Boursiac *et al*., 2010), EICBP.B (or CATMA1; (Huang *et al*., 2019)), all three coding calcium-sensor proteins, are known as negative regulators of SA accumulation and biosynthesis, as well as the light signaling factor FAR1 (Wang *et al*., 2015) or the SA glucosyltransferase UGT74F1 which convert SA in inactive SA 2-O-beta-D-glucoside or the glucose ester of SA (Song *et al*., 2008). On the contrary EDS1, PAD4 (reviewed in (Vlot *et al*., 2009)) and MKS1 (Andreasson *et al*., 2005) are known as positive regulators of SA accumulation. In our data, the repression of CBP60a, ACA11 and EICBP.B genes could sustain a SA biosynthesis and accumulation. In addition, EDS1 activation could allow to consider a positive feedback loop likely to potentiate SA action via EDS1 cytosolic homodimers, even though PAD4 was repressed. But, as well as WRKY70 induction, the repression of the MAPK MKS1 and the activation of the light signaling factor FAR1 (2 times) or the SA glucosyltransferase UGT74F1 could be in favor of less free SA. Concerning SAR, MES1 is known as required in healthy systemic tissues of infected plants to release the active SA from methyl-SA, which serves as a long-distance signal for SAR (Xia *et al*., 2012) and ACBP6 may be involved in the generation of SAR inducing signal(s) (Xia *et al*., 2012). In our data, SAR seemed compromised given the repression of ACBP6 at 24 hpi and MES1 at 72 hpi. To conclude, in the nonhost interaction between pear and *V. inaequalis*, the behavior of some DEGs led us to the hypothesis that the SA pathway could be engaged but transiently and presumably reduced to the few infection sites and not spread by SAR in healthy systemic tissues.

### Calcium influx and reactive oxygen species (ROS) production seems to act as secondary messengers and could lead to stomatal closure in pear / *V. inaequalis* interaction

Early responses of plants upon pathogen perception include calcium influx and ROS production, which both act as secondary messengers (Boller *et al*;, 2009, reviewed in (Lee *et al*., 2017a)). Three pear DEGs were found that seems to indicate early increased cytosolic calcium level. The CSC (Calcium permeable Stress-gated cation Channel) ERD4 (found two times) and the two glutamate receptors GLR3.4 and GLR2.7, are known as calcium permeable channels (Vincill *et al*., 2012, Hou *et al*., 2014). They were induced at 24 hpi in our data. An increased cytosolic calcium level can lead to a pre-invasive defense response by stomatal closure and promote the post-invasive defense response ROS accumulation (Fonseca *et al*., 2019).

Calcium influx has been reported to promote stomatal closure through the regulation of potassium flux and the activation of anion channels in guard cells (reviewed in (Lee *et al*., 2017a)). The stomata closure is known to be induced via the inhibition of inward potassium currents which is achieved via activation of calcium dependent protein kinases (CDPK) such as CPK13 and CPK8/CDPK19 (Ronzier *et al*., 2014, Zou *et al*., 2015); but also via activation of CBL1 of the CBL1-CIPK5 complex, which activates the GORK potassium outward channel (Förster *et al*., 2019). CPK13, CPK8/CDPK19 and CBL1 were all activated at 24 hpi in our data.

A NADPH oxidase RBOHB (respiratory burst oxidase homologs, RBOH) is early (i. e. 24 hpi) and sustainably (through to 72 hpi) induced in the pear/*V. inaequalis* nonhost interaction, which could suggest a rapid and maintained apoplastic ROS production. Indeed, the apoplastic ROS are mainly produced by plasma membrane localized NADPH oxidases, cell wall peroxidases and amine oxidases (kadota *et al*., 2015). In addition, posttranslational regulation of RBOH is required for its activation and ROS production. Calcium, phosphatidic acid, and direct interactors such as Rac1 GTPase and RACK1 (Receptor for Activated C-Kinase 1) have been reported to be positive regulators of RBOHs (reviewed in (Adachi *et al*., 2015)). For example, the Rac-like/ROP GTPase ARAC3 is known to interact with a RBOH to promote ROS production (Zhai *et al*., 2018). In our data, RBOHB activity could also be supported by the presence of positive regulators such as Rac-like/ROP GTPase. The three Rac-like/ROP GTPase ARAC1, ARAC3 and the homolog of At4g03100 were induced at 24 hpi. CDPKs such as CPK1 are also known to activate RBOHs in response to increased cytosolic calcium level (Gao *et al*., 2013). But repression of CPK1 in our data could indicate that this way of activation did not function.

In response to abscisic acid (ABA) or microbe-associated molecular pattern (MAMP) immunity, stomatal closure is known to be regulated by apoplastic ROS production (reviewed in (Qi *et al*., 2017)) and cysteine-rich receptor-like kinases (CRK) are also known to be elements between ROS production and downstream signaling leading to stomatal closure, sometimes activated (CRK10), sometimes inhibited (CRK2 and CRK29; Bourdais *et al*., 2015). Three DEGs coding for CRK were found in our data and the repression of CRK2 and CRK29 (found two times) could be consistent with the stomata closure previously hypothesized, but the repression of CRK10 (found two times) was not. Beyond closure, inhibition of stomatal development could be seen as an extreme defense. YODA (found two times) and MPK6 (found two times) MAPKs belong to a pathway involved in the negative regulation of stomata development (Sun *et al*., 2018). These two genes were induced early in our data.

To conclude, in pear/*V. inaequalis* nonhost interaction, a calcium influx could lead to the development of the stomatal closure pre-invasive defense, but could also promote a post-invasive defense: apoplastic ROS accumulation. Apoplastic ROS, acting themselves as messengers, could come to strengthen the stomatal closure (Fig. 4).

**Figure 4:**
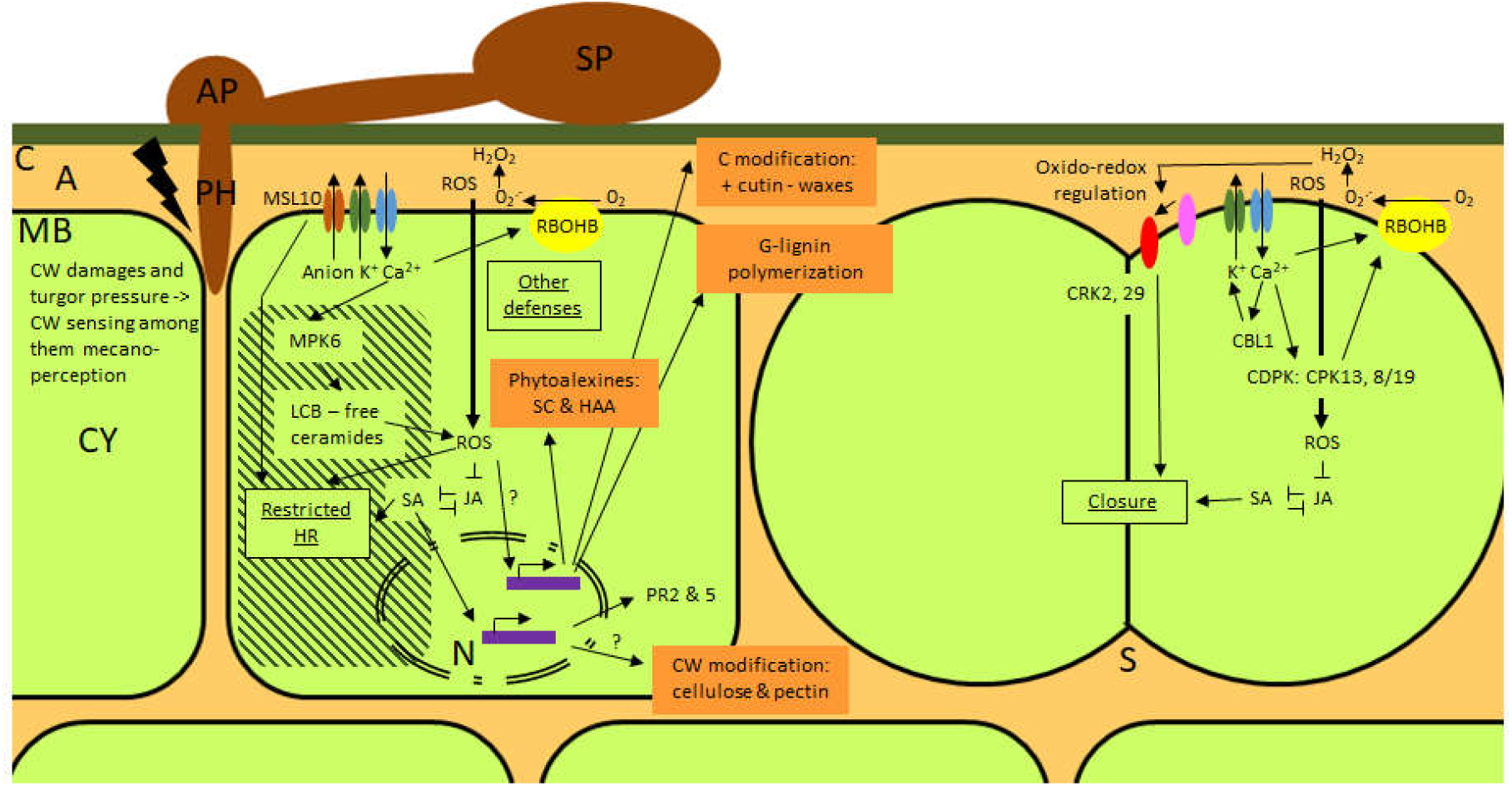
On the left side, events observed in a typical cell, on the right side, events observed in guard cells of a stomata. A: apoplasm, AP: appressorium, C: cuticle, CBL1: calcineurin B-like protein 1, CDPK: Ca2+-dependent protein kinases, CRK: cysteine-rich receptor-like kinase, CY: cytoplasm, CW: cell wall, HAA: hydroxycinnamic acid amines, HR: hypersensitive response, JA: jasmonic acid, MB: plasma membrane, LCB: Long Chain/sphingoid Base components, MPK6: Mitogen activated protein kinase 6, MSL10: mechano-sensitive like 10, N: nucleus, PH: penetration hypha, PR: pathogenesis related proteins, RBOHB: respiratory burst oxidase homolog B, ROS: reactive oxygen species, S: stomata, SA: salycilic acid, SC: simple coumarins, SP: spore

### Transcription factors and sphingolipids could be implicated in maintaining HR under control in pear / *V. inaequalis* interaction

ROS are known to mediate cellular signaling associated with defense-related gene expression, HR i.e. the programmed cell death (PCD) at the site of infection during a pathogen attack, and phytoalexin production (O’Brien *et al*., 2012). Arabidopsis thaliana RCD1 regulator has been proposed to positively regulate cell death in response to apoplastic ROS by protein-protein interactions with transcription factors (reviewed in (Brosché *et al*., 2014)) and WRKY70 and SGT1b were identified as cell death positive regulators functioning downstream of RCD1 (Brosché *et al*., 2014). RCD1 and WRKY70 genes were found induced in our data, at 24 hpi and 72 hpi respectively.

In Arabidopsis, the F-box protein CPR1, in association with the Skp1-Cullin-F-box (SCF) ubiquitin ligase complex, targets for degradation NLR (nucleotide-binding domain and leucine-rich repeats containing proteins) resistance protein such as SNC1, RPM1 or RPS2, to prevent overaccumulation and autoimmunity (reviewed in (Cheng *et al*., 2011)). A Skp1-like (ASK19; 72 hpi) gene and CPR1 (24 hpi) gene were found induced in our data. A gene coding for RPM1 function was also found repressed at 24 hpi. These results could be in favor of the hypothesis that NLR receptors do not take part in the HR development observed in the pear/*V. inaequalis* nonhost interaction (Fig. 1C, H, J). In addition, the induction of an AtSerpin1 gene homolog at 24 hpi (found two times) in our data could be consistent with that hypothesis. Indeed, AtMC1 is a pro-death caspase-like protein required for full HR mediated by intracellular NB-LRR immune receptor proteins such as RPP4 and RPM1 (Coll *et al*., 2010) and AtSerpin1 is a protease inhibitor which block AtMC1 self-processing and inhibit AtMC1-mediated cell death (Lema-Asqui *et al*., 2018).

The differential expression of two others components of the proteasome pathway could be in favor of the HR development: the induction of the RIN3 ubiquitin E3 ligase (24 hpi) and the repression of the BRG3 ubiquitin E3 ligase (24 hpi). Indeed, RIN3 is known as positive regulator of RPM1 dependent HR (Kawasaki *et al*., 2005). And BRG3 is known as a negative regulator of HR in plant/necrotrophic pathogen interactions (Luo *et al*., 2010).

Sphingolipids are involved in the control of PCD, either as structural components of membranes but also as initiators in the cell death regulatory pathway. According to Huby *et al*. (Huby *et al*., 2020), free ceramides and long chain/sphingoid base components (LCBs) are able to trigger cell death, via ROS production, whereas their phosphorylated counterparts, ceramide phosphates and long chain base phosphate components (LCB-Ps) promote cell survival. The induction of PCD by LCB is based on the activation of protein kinases, among them MPK6 (Saucedo-Garcia *et al*., 2011). As already mentioned, MPK6 was found early induced in our data and we found numerous DEGs in the nonhost interaction between pear and *V. inaequalis* that seems to indicate the presence of free ceramides and LCB, which could possibly participate to the HR development. Free LCB presence is suggested by the activations of SBH1 (24 hpi), SLD1 (24 hpi) and another sphingolipid ∆8 long-chain base desaturase homolog to At2g46210 (24 hpi; found two times), and their relative conversion in ceramides is suggested by the differential expressions of the ceramide synthases LOH2 (repressed at 24 hpi) and LOH3/LAG13 (induced at 24 and 72 hpi). LCB non-conversion in phosphorylated counterparts could be shown by the AtLCBK1 repression (72 hpi) and free ceramides maintenance could be suggested by their non-conversion in glycosyled ones given the repression of a glucosyl ceramide synthase homolog to At2g19880 (24 hpi).

The differential expression of numerous known regulators of HR in our data seems again consistent with the HR phenotype observed. The mechanosensor MSL10 and the calmodulin-activated Ca2+ pump (autoinhibited Ca2+-ATPase [ACA]) ACA11 were found differentially expressed: at 24 hpi MSL10 was induced and ACA11 was repressed. MSL10 is known as a positive regulator of cell death (Veley *et al*., 2014) and ACA11 is known as a negative regulator of SA-dependent PCD (Boursiac *et al*., 2010). Their modulation could be linked with the hypothesized calcium influx discussed above (Boursiac *et al*., 2010, Guerringue *et al*., 2018). The participation of the SA pathway in the development of the HR could also be supported by the repression of EDR1 (at 72 hpi). Indeed, the MAPKKK EDR1 is known as a negative regulator of the SA-dependent HR (reviewed in (Zhao *et al*., 2014)).

Three other regulators of HR were found modulated in our data. The transcription factor AS1 (MYB91) was found induced at 24 hpi. It is known as a positive regulator of HR and implicated in the JA pathway (reviewed in [30]). The transcription factor WRKY40 was found repressed at 72 hpi. It is known as a negative regulator of HR (Lee *et al*., 2017b) and implicated in PTI (Najafi *et al*., 2020). Another negative regulator of HR is the lipid-binding domains containing protein VAD1 (Khafif *et al*, 2017). It was found repressed at 72 hpi.

The behavior of another gene in our data, UGT73B3, could indicate that the developed HR was contained due to intracellular ROS production and damages. The function UGT73B3 was thus activated (24 hpi). UGT73B3 is known as a restrictor of HR expansion via its action in detoxification of ROS-reactive secondary metabolites (UGT73B3; (Simon *et al*., 2014)).

To conclude, in pear/*V. inaequalis* nonhost interaction, HR was spread out, potentially in link with a calcium influx and ROS production via free sphingolipids accumulation and not via NLR receptors. Furthermore, the behavior of eight regulators seems to indicate that the developed HR is under control (Fig. 4).

### Cell wall carbohydrates content and cuticle composition could be altered in pear / *V. inaequalis* interaction

The first obstacle encountered by host as well as nonhost pathogens attempting to colonize plant tissues is the plant cell wall, which is often covered with a cuticle. In nonhost resistance, in which non adapted pathogens normally fail to penetrate nonhost plant cells, the cell wall is considered an important factor and seen as a preinvasive penetration barrier, or as the onset place of defensive signaling pathways (Lee *et al*., 2017a, Fonseca *et al*., 2019). Plant cell wall alterations have been demonstrated to have a significant impact on disease resistance and/or on abiotic stresses. These alterations can concern the carbohydrates or the phenolic components, and results in impairing or overexpressing cell wall-related genes (reviewed in (Bacete *et al*., 2018) and (Miedes *et al*., 2014)).

We found numerous genes related to the cell wall with a modified expression during nonhost interaction between pear and *V. inaequalis*, among them about thirty related to the biosynthesis or the modification of carbohydrates. These DEGs are presented in Table 4 except those related to the lignin and other phenolic compounds, which will be discussed later. We saw in particular several DEGs related to cellulose (8) and even more DEGs related to pectin (14) but no DEGs related to callose.

**Table 4:** Main DEGs related to cell wall carbohydrates synthesis/modification detected during nonhost interaction pear/*V. inaequalis*

|  | Gene | Action | Expression* |
| --- | --- | --- | --- |
| Primary cell wall |  |  |  |
| Cellulose | *CSLA2* | synthesis | I |
|  | *PNT1* | synthesis | I |
|  | *COBL2* | deposition (GPI-anchored protein) | R |
|  | *AtGH9A4* | catabolism | I |
|  | *XTR7* | loosening | I |
| Hemi-cellulose (xyloglucan) |  |  |  |
|  | *At5g15490* | synthesis | I |
| Pectin | *At3g42180* | synthesis | I |
|  | *At4g01220* | synthesis | I |
|  | *GHMP kinase* | synthesis | I |
|  | *RHM1* | synthesis | R |
|  | *PME At2g45220* | methylesterification | I |
|  | *PME At2g46930* | methylesterification | I |
|  | *PME At3g05910* | methylesterification | R |
|  | *PME At1g02810* | methylesterification | R |
|  | *PME44* | methylesterification | R |
|  | *PG At3g16850* | depolymerisation | I |
|  | *PG At3g59850* | depolymerisation | I |
|  | *PG At4g13710* | depolymerisation | I |
|  | *PG At3g62110* | depolymerisation | R |
|  | *IDA* | degradation | R |
| Arabinogalactan protein |  |  |  |
|  | *AGP11* | – | I |
|  | *AGP1* | – | R |
| Secondary cell wall |  |  |  |
| Cellulose | *CESA09* | synthesis | I |
|  | *CESA10* | synthesis | I |
|  | *CSLG1* | synthesis | I |
| Hemi-cellulose (xylan) | *FRA8* | synthesis | I |
| Undetermined |  |  |  |
| Expansin | *EXP15* | loosening | I |
|  | *EXPB3* | loosening | I |
| Hemi-cellulose | *ATFUC1* | modification | I |
|  | *XTH33* | growth and assembling | R |

Note: \*I: induced, R: repressed

Concerning these particular carbohydrate components, the model proposed by Bacete *et al*. (Bacete *et al*., 2018) is as follows. Firstly, alterations in cellulose biosynthesis from primary or secondary cell wall trigger specific defensive responses, such as those mediated by the hormones JA, ET or abscisic acid (ABA), activate biosynthesis of antimicrobial compounds, but also might attenuate pattern triggered immunity (PTI) responses. Secondly, alterations of cell wall pectins, either in their overall content, their degree of acetylation or methylation, activate specific defensive responses, such as those regulated by JA or SA, and trigger PTI responses, probably mediated by damage-associated molecular patterns like oligogalacturonides. Thus, even though our results do not completely support a role of these DEGs, we suppose that the modified expression of cell wall related genes during nonhost interaction between pear and *V. inaequalis* could be meaningful.

Concerning the cuticle layer, most cuticles are composed largely of cutin, an insoluble polyester of primarily long-chain hydroxy fatty acids. This lipophilic cutin framework is associated with hydrophobic compounds collectively referred to as waxes. The cuticle is also thought to contain varying amounts of intermingled cell wall polysaccharides and sometimes also a fraction termed cutan (reviewed in (Fich *et al*., 2016)). Cutin monomers are synthesized by the modification of plastid-derived 16C and 18C fatty acids in the ER, yielding variously oxygenated fatty acid–glycerol esters referred to as monoacylglycerols, which polymerize upon arrival at the growing cuticle (Fig. 5, reviewed in (Fich *et al*., 2016)).

**Figure 5:**
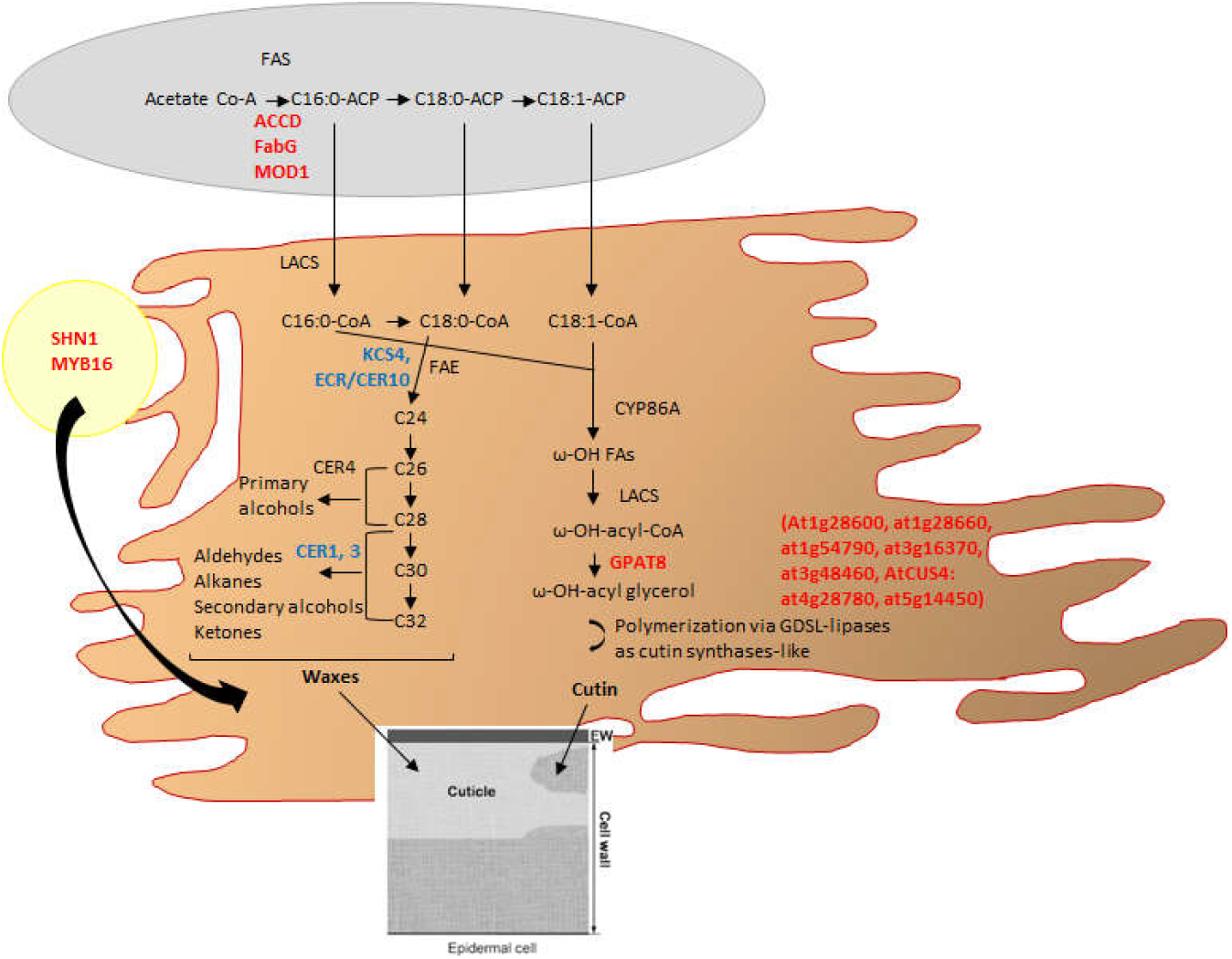
In green the chloroplast, in brown the ER and in yellow the nucleus. Genes written in red are induced, genes written in blue are repressed. FAS: Fatty Acid Synthase complex to which belong ACCD (carboxytransferase beta subunit of the Acetyl-CoA carboxylase complex), FabG (β-ketoacyl ACP-reductase) and MOD1 (enoyl-ACP-reductase) functions. FAE: fatty acid elongase complex. KCS4 (3-ketoacyl-CoA synthase 4) and ECR/CER10 (trans-2-enoyl-CoA reductase) belong to the FAE complex. CER1 (octadecanal decarbonylase) and CER3 are implicated in aldehydes (CER1) and alkanes (CER1 and 3) generation in waxes biosynthesis. In cutin monomers synthesis, the ω-hydroxylation of C16:0 and C18:1 is catalyzed by cytochrome P450 monooxygenase (CYP86A) and LACS-encoded acyl-CoA synthetase may be required either to synthesize 16-hydroxy 16:0-CoA, a substrate for ω-hydroxylase, or for membrane transfer of monomers. Finally, the mature monoacylglycerol cutin monomers are generated by transfer of the acyl group from acyl-CoA to glycerol-3-phosphate by glycerol-3-phosphate acyltransferase (GPAT) enzymes such as GPAT8. Some GDSL-lipases enzyme (such as At1g28600, At1g28660, At1g54790, At3g16370, At3g48460, AtCUS4: At4g28780, At5g14450) are then functioning as cutin synthase and polymerize cutin monoacylglycerols. Transcription factors such as MYB16 and SHN1 are positive regulators of wax and cutin biosynthesis. Adapted from Fich *et al*., 2016 and Lim *et al*., 2017

C16 and C18 fatty acids are also important precursors of cuticular wax synthesis (Fig. 5). Upon transport to the ER, the C16 and C18 fatty acids are extended to form very-long-chain fatty acids (VCLFAs; C>20), and this extension is carried out by the fatty acid elongase (FAE) complex located on the ER membrane. The very-long-chain FAs are then converted into the varied cuticular waxes (primary alcohols, aldehydes, alkanes, secondary alcohols, ketones) by many ways (reviewed in (Lim *et al*., 2017)).

Interestingly, we found three genes up-regulated 24 hpi belonging to the FAS (fatty acid synthase) chloroplastic complex implicated in the production of the C16 precursor (Fig. 5): ACCD, FabG and MOD1 (found two times). ACCD encodes the carboxytransferase beta subunit of the Acetyl-CoA carboxylase complex which catalyzes the first committed step in fatty acid synthesis: the carboxylation of acetyl-CoA to produce malonyl-CoA. FabG and MOD1 are respectively a β-ketoacyl ACP-reductase and an enoyl-ACP-reductase which catalyze respectively the conversion of acetoacetyl-ACP into β-hydroxyacyl-ACP and the second reductive step from enoyl-ACP to butyryl-ACP (reviewed in (Lim *et al*., 2017)).

In the ER, the four functions we found related to waxes biosynthesis in our data were repressed at 24 hpi: KCS4 (found two times), CER1 and CER3, or 72 hpi: ECR/CER10. KCS4 and ECR/CER10 belong to the FAE complex (Joubès *et al*., 2008; Lee *et al*., 2015). The last two genes are implicated in aldehydes (CER1) and alkanes (CER1 and 3) generation (reviewed in (Lim *et al*., 2017)). On the contrary, the eight genes we found related to cutin biosynthesis were induced at 24 hpi except a gene homolog to At5g14450, which was induced at 72 hpi. One of them is a glycerol-3-phosphate acyltransferase (GPAT) coding gene: GPAT8, which catalyzes the transfer of a fatty acid from coenzyme A (CoA) to glycerol-3-phosphate (Fig. 4; reviewed in (Fich *et al*., 2016)). GPAT8 function in cutin formation has been functionally confirmed in association with GPAT4 (Li *et al*., 2007). The seven others genes code GDSL-lipases enzyme (At1g28600, At1g28660, At1g54790, At3g16370, At3g48460, AtCUS4: At4g28780, At5g14450), some of which have been shown to function as cutin synthase (Fig. 4; Yeats *et al*., 2014; reviewed in (Fich *et al*., 2016)) and polymerize monoacylglycerols.

We also found induced respectively at 24 and 72 hpi two genes involved in waxes and cutin biosynthesis positive regulation: MYB16 and SHN1. The SHN genes (SHN1–SHN3), a set of three largely redundant APETALA 2 family transcription factors from A. thaliana, are regulators of floral cutin and epidermal cell morphology. SHN1 is regulated by the MYB family transcription factor MYB106, which, along with its paralog MYB16, controls many aspects of cuticle and epidermis formation in A. thaliana (reviewed in (Cui *et al*., 2016) and (Fich *et al*., 2016)).

Cutin and cuticular waxes play an important role in plant-insect and plant-microbe interactions. Numerous Arabidopsis mutants in cutin and waxes biosynthetic or transport genes, such as Acyl-CoA binding proteins (ACBP), show varying degrees of cuticle impairment, alterations in cutin and/or wax composition, and defects in SAR (reviewed in (Lim *et al*., 2017). We found ACBP6 repressed at 24 hpi. That repression is not inconsistent with the previously described possible amplification of cutin biosynthesis and polymerization, given that acbp6 KO mutation is not associated with a defect in that pathway (Xia *et al*., 2012). That repression is also consistent with the SAR repression hypothesized above as the acbp6 KO mutant show compromised SAR (Xia *et al*., 2012).

To conclude, our analysis of nonhost pear/*V. inaequalis* interaction could show an alteration of the cuticle composition with more cutin and less waxes synthesis. The potential increase in cutin polymerization could lead to a thickening of the cuticular layer to prevent fungus penetration via its appressoria.

### Secondary metabolism seems to lead to G unit lignin polymerization and simple coumarin or hydroxycinnamic acid amine phytoalexins synthesis in pear / *V. inaequalis* interaction

As distinguished from primary metabolism, plant secondary metabolism refers to pathways and small molecule products of metabolism that are non-essential for the survival of the organism. But they are key components for plants to interact with the environment in the adaptation to both biotic and abiotic stress conditions. Plant secondary metabolites are usually classified according to their chemical structure. Several groups of large molecules, including phenolic acids and flavonoids, terpenoids and steroids, and alkaloids have been implicated in the activation and reinforcement of defense mechanisms in plants (reviewed in (Yang *et al*., 2018)).

Terpenoids and steroids, or isoprenoids, are components of both the primary and secondary metabolisms in cells, and mono-, tri-, sesqui- and polyterpenes are considered as secondary metabolites (reviewed in (Tetali *et al*., 2019)). Our results on pear identified seven DEGs and five DEGs which could belong to the chloroplastic methylerythritol posphate (MEP) and to the cytosolic mevalonic acid (MVA) pathway of isoprenoids production respectively (Table 5), which results, among others compounds, in tri- and sesquiterpenes secondary metabolites. The majority of these genes contribute to produce primary metabolites according to Tetali (Tetali *et al*;, 2019). Except SMT2, that we found induced at 24 hpi, there is no report concerning a putative implication of others genes in plant biotic resistance. SMT2 is involved in sterols production and smt2 mutation was reported to compromise bacterial resistance in Nicotiana benthamiana (Wang *et al*., 2012). The hypothesis is that sterols regulate plant innate immunity against bacterial host and nonhost infections by regulating nutrient efflux into the apoplast. *V. inaequalis* is a hemibiotrophic pathogen which colonizes only the apoplast compartment at the beginning of the interaction. Strong relative induction of SMT2 in our data could indicate that a similar mechanism of nutrient efflux regulation via sterols could take place to limit the fungus growth in pear nonhost resistance against *V. inaequalis*.

**Table 5:**
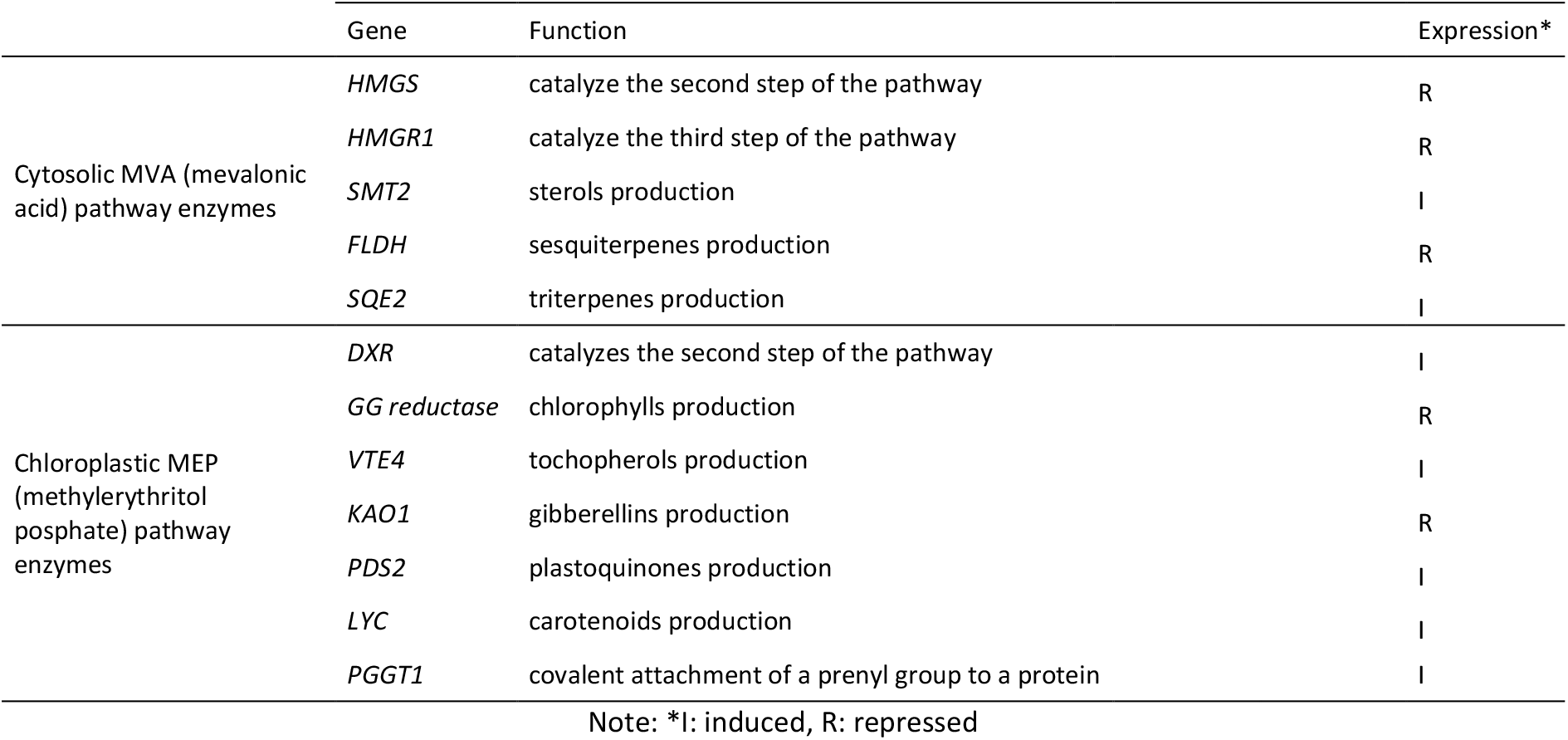
Main DEGs involved in biosynthetic pathways for terpenes and isoprenoids during pear/*V. inaequalis* nonhost interaction

In our data, the other DEGs that were linked to secondary metabolism belong to the phenylpropanoid pathway production (Fig. 6). Among them we found four genes which could belong to the flavonoid production, all repressed, at 24 hpi (DFR and DRM6) or 72 hpi (TT7 and UGT71D1). DFR (dihydroflavonol reductase) is involved in flavan-3,4-ol production and TT7 (flavonoid 3’ hydroxylase) in dihydroquercetin production from dihydro-kaempferol, and UGT71D1 (glucosyl transferase) in quercetin-glycoside production from quercetin (TAIR database; https://www.arabidopsis.org/index.jsp). DMR6 (flavone synthase) is involved in flavone production from naringenin (Falcone *et al*., 2015). Thus flavonoid production does not seem to be favored, which is not consistent with the induction of MYB12 at 24 hpi, but consistent with MYB4 induction at 72 hpi. MYB12 is actually known as a positive regulator of flavonol biosynthesis in pear and apple fruits (Wang *et al*., 2017b, Zhai *et al*., 2019) whereas MYB4 is known as a negative regulator of this biosynthetic pathway (Wang *et al*., 2020).

**Figure 6:**
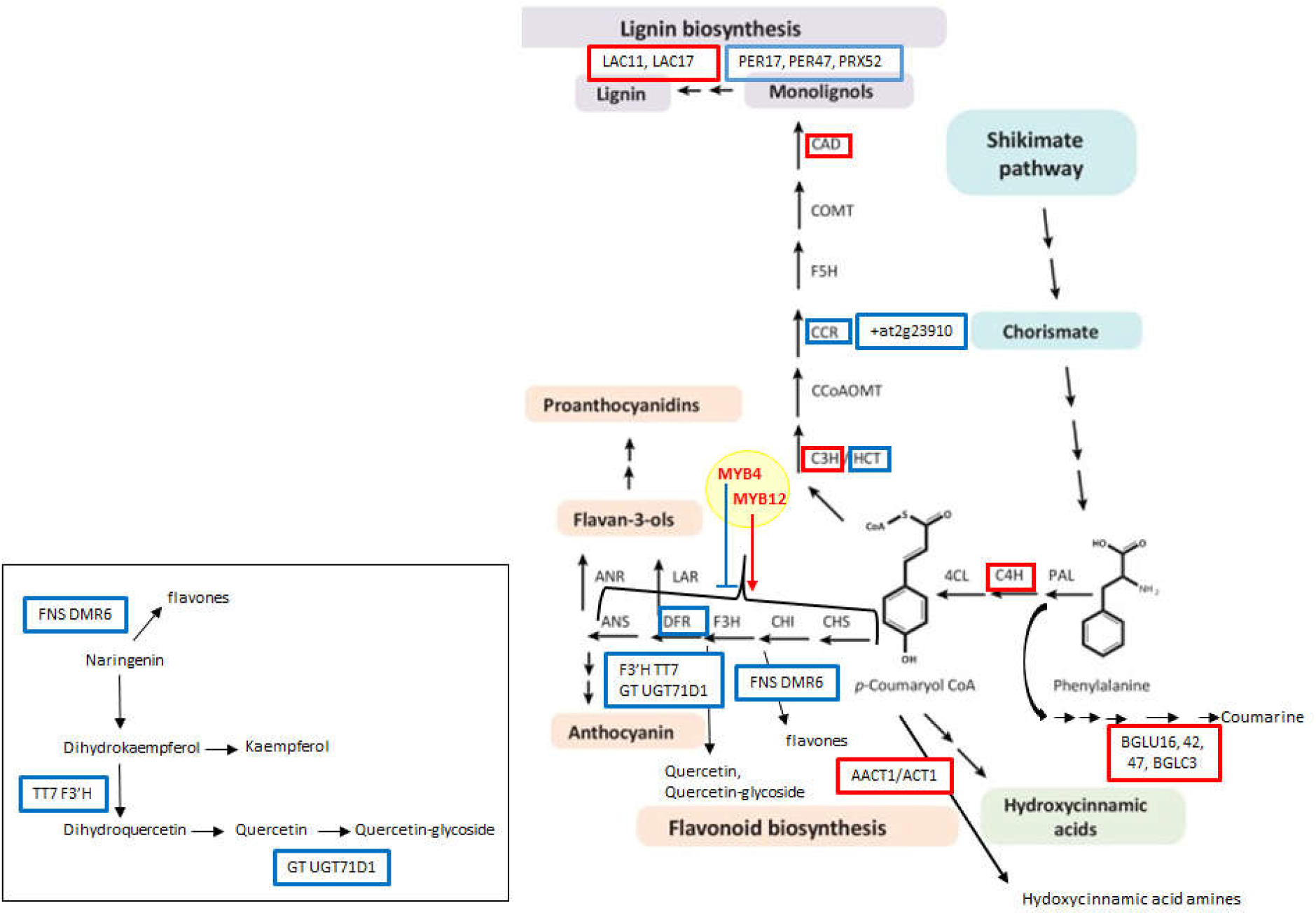
Genes framed in red are induced, genes frames in blue are repressed. Framed in black, the detail of genes involved in flavonoids production and found in this interaction. Abbreviations: 4CL, 4-coumarate-CoA ligase; AACT, anthocyanin 5-aromatic acyltransferase; ANR, anthocyanidin reductase; ANS, anthocyanin synthase; BGLC or BGLU, β-glucosidases; C3H, coumarate 3-hydroxylase; C4H, cinnamate 4-hydroxylase; CAD, cinnamyl alcohol dehydrogenase; CCoAOMT, caffeoyl-CoA O-methyltransferase; CCR, cinnamoyl-CoA reductase; CHI, chalcone isomerase; CHS, chalcone synthase; COMT, caffeic acid 3-O-methyltransferase; CPK, calcium-dependent protein kinase ; DFR, dihydroflavonol reductase; DMR6, downy mildiou resistant 6; F3H, flavanone 3-hydroxylase; F3’H flavonoid 3’-hydroxylase; FLS, flavonol synthase; FNS, flavone synthase; GGT1, gamma-glutamyl transpeptidase 1; GT, glucosyl transferase; HCT, hydroxycinnamoyl-CoA shikimate/quinate hydroxycinnamoyl transferase; LAC, laccase; LAR, leucoanthocyanidin reductase; OMT1, O-methyltransferase 1; PAL, phenylalanine ammonia-lyase; PER or PRX, peroxidase; TT7, transparent testa 7; UGFT, UDP-glucose flavonoid-3-O-glucosyltransferase; UGT71D1, UDP-glycosyltransferase 71D1

Concerning the production of monolignols, precursors of lignin synthesis, some genes potentially related were found induced, others repressed. We found CYP98A3 and CAD9 (found two times) induced at 24 hpi and HCT, CCR1 and a gene homolog to At2g23910 (found two times, one time repressed at 24 hpi, one time repressed at 72 hpi, Fig. 6). CYP98A3 encodes a C3H (coumarate 3-hydroxylase), CAD9 encodes a CAD (cinnamyl alcohol dehydrogenase), HCT is an hydroxycinnamoyl-CoA shikimate/quinate hydroxycinnamoyl transferase, CCR1 encodes a CCR (cinnamoyl-CoA reductase) and At2g23910 encodes a CCR-related protein. (TAIR and KEGG databases (https://www.genome.jp/kegg/)).

Lignification is obtained by cross-linking reactions of the lignin monomers or by polymer–polymer coupling via radicals produced by oxidases such as peroxidases (Fernandez-Perez *et al*., 2015a) and laccases (Zhao *et al*., 2013). However, while peroxidases are able to oxidize monolignols to produce H, G and S units of lignin, laccases only generate G units (Fernandez-Perez *et al*., 2015a). In our data, we found two laccases induced at 24 hpi: LAC11 (found two times, one time induced at 24 and 72 hpi) and LAC17 (found two times), and three peroxidases repressed at 24 hpi: PRX17, PER47 and PRX52 (also repressed at 72 hpi), which could be linked to lignin biosynthetic process (Fig. 6). According to Zhao *et al*. (Zhao *et al*., 2013), LAC11 and LAC17, along with LAC4, play a critical role in lignification, and their results suggests that peroxidase and laccase do not serve redundant functions in lignification, at least in the vascular tissues of the stem and root of Arabidopsis. Participation in lignin formation has also been proved for PRX17 (Cosio *et al*., 2017), PER47 (Tokunaga *et al*., 2009) and PRX52 (Fernandez-Perez *et al*., 2015b). But there are currently no reports about a possible involvement of all these genes in lignification linked to biotic or abiotic stresses. Concerning nonhost resistance, two reports describe lignin accumulation/deposition involvement: one in apple fruit (Vilanova *et al*., 2012) and the other one in cowpea (Fink *et al*., 1991). In the latest, authors showed that preferentially generated lignin units in this nonhost interaction are G units, just as it could be the case in our pear / *V. inaequalis* study. To summarize, it is tempting to think that modifications of expression observed for genes linked to lignin polymerization are relevant for the pear nonhost resistance against *V. inaequalis*, but further functional analysis should be conducted to conclude about the lignin status, such as histochemical staining (Yu *et al*., 2022), content measure by absorbance (Yu *et al*., 2022) or Fourier-transform infrared (FTIR) spectroscopy analysis (Ranade *et al*., 2022).

The biosynthesis of two others types of phenylpropanoid compounds could appear to be favored during pear nonhost resistance against *V. inaequalis*: simple coumarin on one hand and hydroxycinnamic acid amides on the other hand. We found four BGLU-like genes induced: BGLU47 and BGLC3 (at 24 hpi), BGLU16 (at 72 hpi); BGLU42 (at 24 and 72 hpi) (Fig. 6). These β-glucosidases could be implied in simple coumarin path production from the cinnamic acid (KEGG database). Some natural simple coumarins are known as antifungal compounds in vitro and have been developed as fungicides (Song *et al*., 2017). Previous work on Hevea also reports the correlation between the resistance against pathogenic fungi and the production of some coumarins, with antifungal activity in vitro (Guisemann *et al*., 1986). We also found induced at 24 hpi the genes AACT1/ACT1, ATPAO5 and genes homologs to At4g17830 and At4g38220 (Fig. 6). AACT1/ACT1 catalyze the first specific step in branch pathway synthesizing hydroxycinnamic acid amides from the p-Coumaroyl CoA or the feruloyl CoA and amines agmatine or putrescine (Muroi *et al*., 2009). Hydroxycinnamic acid amides are produced in response to pathogenic infections (Muroi *et al*., 2009) and surface exported. Hydroxycinnamic acid amides are reported to participate in Arabidopsis nonhost resistance against Phytophthora infestans via their inhibitory activity on spore germination (Dobritzsch *et al*., 2016). The three others genes belong to the arginine biosynthesis path (homologs to At4g1783 and At4g38220) and the arginine and proline metabolisms which produce the amines agmatine and putrescine (ATPAO5) (KEGG database). Agmatine is directly produced from arginine thanks to an ADC activity (arginine decarboxylase) and putrescine can be produced from spermidine thanks to a PAO activity (polyamine oxidase). ATPAO5 catalyzes the conversion of spermine in spermidine. The induction of these three last genes is therefore consistent with the hypothesis of amines production in order to enable hydroxycinnamic acid amides synthesis. The induction of C4H at 24 hpi could also favor hydroxycinnamic acid amides synthesis via p-Coumaroyl CoA biosynthesis promotion. C4H (cinnamate 4-hydroxylase) catalyzes the production of p-Coumaric acid from Cinnamic acid and p-Coumaric acid gives p-Coumaroyl CoA thanks to 4CL (4-coumarate-CoA ligase) (KEGG database).

Among the suite of defense components synthetized in nonhost as in host context, a chemical barrier can be established via accumulation of a diverse array of secondary metabolites rapidly produced upon pathogen infection, named phytoalexins, with toxic or inhibitory effects (reviewed in (Lee *et al*., 2017a)). Phytoalexins can be flavonoids, such as the pisatin of pea (in Celoy *et al*., 2014) but also varied phenylpropanoid compounds. In the nonhost interaction pear / *V. inaequalis*, the production of flavonoid type phytoalexins does not seem to be favored, except possibly simple coumarin and hydroxicinnamic acid amines.

### Very limited transcriptomic modulation during apple / *V. pyrina* nonhost interaction

Only 60 DEGs were detected in the apple / *V. pyrina* nonhost interaction at 24 or 72 hpi, in agreement with the total absence of macroscopic symptoms and few cells engaged in an HR-like reaction observed at the microscopic level. Among these 60 DEGs, 36 have unpredicted function. Among the 24 remaining DEGs, nine DEGS could be relevant in apple / *V. pyrina* nonhost interaction in view of our findings in pear / *V. inaequalis* nonhost interaction. ORG2 (BHLH038), a putative integrator of various stress reactions (Vorwieger *et al*., 2007) was induced at 24 hpi. Three genes were related to an oxidative stress: GASA10 was repressed at 24 hpi and NRAMP3 and AOR were induced at 24 hpi. GASA proteins have been suggested to regulate redox homeostasis via restricting the levels of OH. in the cell wall (Trapalis *et al*., 2017). The repression of this gene could be thus in favor of more OH. in the cell wall. The oxidoreductase coding gene AOR is known in the chloroplast to contribute to the detoxification of reactive carbonyls produced under oxidative stress (Yamauchi *et al*., 2012). NRAMP genes function as positive regulators of ROS accumulation, especially during Arabidopsis Erwinia chrisanthemi resistance (Segond *et al*., 2009). The induction (at 24 and 72 hpi) of another gene could suggest modifications at the cell wall level: EXP8, an expansin coding gene involved in cell wall loosening (Tair database). We also found two genes related to hormone pathways, one induced at 24 hpi: WIN1 and the other one repressed at 72 hpi: UBP12. WIN1 is known as a negative regulator of SA pathway (Lee *et al*., 2008) and UBP12 as a positive regulator of JA pathway via the stabilization of MYC2 (Jeong *et al*., 2017). In possible link with the JA pathway, we also found TPS21 induced at 24 hpi. TPS21 is involved in sesquiterpenes production and is promoted by JA signal via MYC2 (Hong *et al*., 2012). TPS21 is especially involved in the jasmonate-dependent defensive emission of herbivore-induced volatiles in cranberries (Rodriguez-Saona *et al*., 2013). Finaly the last DEG we found potentially relevant in apple / *V. pyrina* nonhost interaction could promote HR via ceramides accumulation. ACD11 is repressed at 24 hpi in our data. In acd11 mutants, the relatively abundant cell death inducer phytoceramide rises acutely (Simanshu *et al*., 2014).

Because nonhost resistance of apple against *V. pyrina* has resulted in a very limited number of cells engaged in an HR-like reaction, it has not been possible for us to exhaustively describe its outcome at the transcriptomic level in the first three days of the interaction. Further insight with later points of kinetic and more adapted techniques such as laser-assisted cell picking, prior to micro arrays or RNA sequencing analysis (review in (Fink *et al*., 2006)) could provide more information in the future.

### Comparison of pear resistances against the host pathogen *V. pyrina* and the nonhost pathogen *V. inaequalis*

Perchepied *et al*. (Perchepied *et al*., 2021) performed a detailed transcriptomic analysis of the host resistance of pear against *V. pyrina* strain VP102, deployed in a transgenic pear bearing the well-known apple Rvi6 resistance gene against *V. inaequalis*. Comparing this work to ours gives us the rare opportunity to analyze similarities and differences between a host and a nonhost resistance in the same plant.

Concerning the recognition and early signaling steps of the interactions, many receptors and co-receptors have been found induced in the host pear resistance, especially damage-associated molecular patterns receptors such as RLK7, revealing that PTI and ETI must be engaged. We did not find evidence of the mobilization of such receptors in the pear nonhost resistance. PTI and ETI receptors are nonetheless reported as implicated in nonhost resistance (reviewed in (Gill *et al*., 2015) and (Lee *et al*., 2017a)). As we only analyzed post infection transcriptional modulations in the nonhost pear/*V. inaequalis* interaction (at 24 and 72 hpi), one hypothesis to explain the lack of PTI and ETI receptors in our data could be that these receptors were already present as preformed defenses and not particularly induced by the infection onset. In pear nonhost interaction, the earliest likely signaling pathways we were able to highlight are calcium influx and apoplastic ROS production. Calcium signaling seems to be also implicated in pear host resistance, but less obviously than it seems in nonhost resistance.

Regarding the hormonal signaling pathways, the JA defense signaling pathway seems to be repressed in pear nonhost resistance but was found quite activated in pear host resistance. The JA/ET defense signaling pathway is known as an effective defense against necrotrophic fungi in Arabidopsis (Pieterse *et al*., 2012). Thus, it is not inconsistent to find the JA pathway potentially repressed in the development of the pear nonhost resistance against the hemi-biotrophic pathogen *V. inaequalis*. But it is very interesting to find this pathway rather induced in the development of the pear host resistance against the other hemi-biotrophic pathogen *V. pyrina*. The SA signaling pathway is commonly seen as the classical one triggered to resist biotrophic fungi in Arabidopsis (Pieterse *et al*., 2012), but it seems only a little engaged in pear nonhost resistance, and is repressed in pear host resistance. If this absence of SA implication is quite unexpected in pear host resistance against a hemi-biotrophic fungus, it could be consistent with the report that the exact role of these key defense phytohormones is unclear in nonhost resistance and remains to be established (Fonseca *et al*., 2019). As shown by Tsuda *et al*. (Tsuda *et al*., 2009), an explanation for the hormone pathways behavior in pear host resistance could be that: as both the SA and JA/ET pathways positively contribute to immunity, a loss of signaling flow through the SA pathway can be compensated by a rerouting signal through the JA/ET pathways. In addition, independently of SA signaling, but in positive connection with JA signaling, SAR seems to be engaged in distal tissues during pear host resistance. To conclude, in pear host as well as nonhost resistances, classical resistance hormones SA and JA/ET, and the correlative PR gene defenses, seems differently involved than in Arabidopsis.

The carbohydrate content of the cell-wall seems to be modified in response to the attacks by the pathogens. Regarding cell-wall and cuticle, in pear host as well as nonhost resistances, the numerous DEGs found could highlight important modifications. Similar modifications affected the cellulose and mainly the pectin contents, but no callose production was observed. Regarding cuticle, waxes production was induced in host resistance whereas it seems to be repressed in nonhost resistance, in favor of cutin production / polymerization, which was also induced in host resistance. To conclude, as a first obstacle encountered by host, as well as nonhost pathogens attempting to colonize plant tissues, the plant cell wall and its cuticle seem to play a leading role in both pear host and nonhost resistances.

Finally, the production of secondary metabolites and phenylpropanoids compounds in particular, could be a major line of defense, in pear host as well as nonhost resistances, but with divergences. If lignin and flavonoid productions are preponderant in pear host resistance against *V. pyrina*, lignin implication in pear nonhost resistance seems less clear and flavonoids production seems repressed. But the biosynthesis of two other types of phenylpropanoid-derived phytoalexins could be favored during pear nonhost resistance: simple coumarin on one hand and hydroxycinnamic acid amides on the other hand.

The comparative analysis between a host and a nonhost resistance in pear shows that, even though specificities are observed, two major defense lines engaged seems shared: the cell wall and its cuticle on one hand, the secondary metabolism with the phenylpropanoid pathway on the other hand. Moreover, these defenses seem deployed independently of the SA signaling pathway, yet widely recognized as the main defense hormone against biotrophic pathogens.

## Conclusion

To our knowledge, our work is the first one published regarding a transcriptomic analysis of post-infections events of a nonhost resistance to *Venturia* sp. in apple and pear. Here, our molecular work on apple / *V. pyrina* nonhost resistance remains preliminary and in order to allow a deeper deciphering, further analyses must be considered with the aid of tools adapted to this nonhost resistance with very few cells engaged in an HR-like reaction, only visible at a microscopic level. In pear, this deciphering allowed us to show that nonhost resistance against *V. inaequalis* involves enough pathogen penetration in plant tissue to occasionally trigger visible HR, and develops post-invasive defenses.

To summarize our findings on pear with a notion of cascading effect, we can propose the following scenario (Fig. 4): once *V. inaequalis* presence is recognized by pear, a calcium cellular influx could be induced and could lead to the possible development of a pre-invasive defense, the stomatal closure, but could also promote a possible early post-invasive defense, an apoplastic ROS accumulation. Apoplastic ROS, acting themselves as ubiquitous messengers, could come to reinforce the stomatal closure but could also mediate cellular signaling possibly resulting in two post-invasive defenses: HR development at infection sites, along with phytoalexin (simple coumarin and hydroxicinnamic acid amines) production. The inferred alterations of the epidermis composition (cellulose, pectin, lignin for the cell wall, and cutin for the cuticle), are presumed to strengthen this physical barrier and could be seen as the development of another pre-invasive defense. Nonhost resistance is defined as the resistance of an entire plant species against a specific parasite or pathogen and is seen as the most durable resistance of plant. Thus, understanding the molecular mechanisms underlying nonhost resistance can open up some interesting avenues to create sustainable host resistances in the same plant species. Considering pear, in order to stop the germination and entrance of hemibiotrophic host fungi such as *V. pyrina*, strengthening the cuticle initial barrier via more cutin production and cross-link, or promoting the biosynthesis of phytoalexins like hydroxycinnamic acid amines, appear as promising solutions.

## Abbreviations

ABA: abscisic acid
CDPK: calcium dependent protein kinase
CRK: cysteine-rich receptor-like kinase
DEG: differentially expressed gene
DFR: dihydroflavonol 4-reductase
ET: ethylene
ER: endoplasmic reticulum
ETI: effector triggered immunity
FAE: fatty acid elongase
GPAT: glycerol-3-phosphate acyltransferase
hpi: hours post inoculation
HR: hypersensitive reaction
JA: jasmonic acid
LCB: long chain/sphingoid base component
LCB-Ps: long chain base phosphate component
LOX: lipoxygenase
MAMP: microbe-associated molecular pattern
OPDA: 12-oxo-phytodienoic acid
PCD: programed cell death
PTI: pattern triggered immunity
RBOH: respiratory burst oxidase homolog
ROS: reactive oxygen species
SA: salicylic acid
SAR: systemic acquired resistance

## Acknowledgements

The authors gratefully acknowledge the IRHS-ImHorPhen team of INRA Angers for technical assistance in plant maintenance and the technical platforms ANAN and IMAC. Preprint version 4 of this article has been peer-reviewed and recommended by Peer Community In Genomics (https://doi.org/10.24072/pci.genomics.100025).

## Data, scripts, code, and supplementary information availability

The datasets supporting the conclusion of this article are available in the Gene Expression Omnibus (GEO) repository [https://www.ncbi.nlm.nih.gov/geo/] with GSE159179 and GSE159180 accession numbers for apple and pear respectively (Vergne, 2020a, Vergne, 2020b). The pipeline AnaDiff used for differential analyses is available at https://doi.org/10.5281/zenodo.6477917 (Pelletier, 2022). Additional information is available in Additional File 1: Table S1, S2, S3, S4 and S5.

